# Mapping energy landscapes of homopolymeric RNAs via simulated tempering and deep unsupervised learning

**DOI:** 10.1101/2023.10.10.561750

**Authors:** Vysakh Ramachandran, Davit A Potoyan

## Abstract

Conformational dynamics plays crucial roles in RNA functions about sensing and responding to environmental signals. The liquid-liquid phase separation of RNAs and the formation of stress granules partly relies on RNA’s conformational plasticity and its ability to engage in multivalent interactions. Recent experiments with homopolymeric and low-complexity RNAs have revealed significant differences in phase separations due to differences in base chemistry of RNA units. We hypothesize that differences in RNA phase-transition dynamics can be traced back to the differences in conformational dynamics of single RNA chains. In the present contribution, we utilize atomistic simulations with numerous unsupervised learning to map temperature dependence conformational free energy landscapes for homopolymeric RNA chains. These landscapes reveal a variety of metastable excited states influenced by the nature of base chemistry. We shed light on the distinct contributions of the polyphosphate backbone versus base chemistry in shaping conformational ensembles of different RNAs. We demonstrate that the experimentally observed temperature-driven shifts in metastable state populations align with experimental phase diagrams for homopolymeric RNAs. The work establishes a microscopic framework to reason about base-specific RNA propensity for phase separation. We believe our work will be valuable for designing novel RNA sensors for biological and synthetic applications.

## INTRODUCTION

Molecules of RNA play crucial roles in nearly every facet of cellular information processing, encompassing catalysis, metabolism, and transcriptional regulation(1–5). The multifunctionality of RNAs is harnessed in both medical and synthetic realms, making RNA molecules convenient as therapeutic targets for genetic diseases and as foundational elements in synthetic biology applications (5). Despite their capability to fold into intricate 3D structures through hierarchical secondary and tertiary formations, RNAs often retain a notable degree of conformational disorder. An emerging perspective on RNA functionality underscores the significance of dynamic conformational ensembles which are precisely calibrated to detect environmental cues such as small molecules, ion atmosphere variations, and temperature fluctuations(6–10).

While homopolymeric RNA sequences appear straightforward in composition, their biological roles are remarkably varied and intricate. Homo ribo-polynucleotide tracts like Poly G, Poly A, Poly C, and Poly U serve diverse functions(11–14). For example, mRNA molecules’ elongated poly(A) tails enhance translation and bolster mRNA stability (15, 16). Similarly, such tracts are found in various non-coding RNAs – like U tracts in long non-coding RNAs (lncRNA) – executing distinct roles. Yet, our understanding of the exact molecular mechanisms underpinning these tracts’ functionalities is limited, largely due to an incomplete understanding of their structural and dynamic features.

A new functional link that may shed light on the importance of long stretches of homopolymeric RNA and low complexity RNA sequence, in general, is the unique ability of RNA to engage in multivalent interactions with other RNAs and proteins, which lead to phase-separation and formation of membraneless organelles such as stress granules and ribonucleoproteins bodies (17, 18). Experiments have shown that homopolymeric RNAs can phase-separate with lower critical solution temperatures (LCSTs)(17–19). Notably, the thermodynamics of this phase separation is inherently sequence-dependent, adhering to a thermal stability hierarchy: Poly(G) > Poly(A) > Poly(C) > Poly(U)(18). Also the recent computational work employing coarse-grained models for RNAs have found conformational transitions between hairpin and extended forms take place even within condensate droplets(20, 21). To fully grasp RNA’s inherent tendency for self-assembly and condensate formation, however, a detailed atomistic characterization of RNA conformational free energy landscape, encompassing an array of metastable structures, is imperative.

In this study, we employed simulated tempering to sample conformations of explicit solvated 20-nucleotide single chains of Poly(G), Poly(A), Poly(C), and Poly(U) chains. We used all backbone dihedral angles of RNA to map free energy landscapes and characterize the main conformational states of RNA chains. To ascertain robustness of conformational landscapes and their features we used three dimensionality reduction techniques; principal component analysis (PCA), time-lagged independent component analysis (TICA), and variational Autoencoders (VAE). Furthermore, we have explored how the free energy landscapes change with temperature which sheds light on the ability of RNA’s to sense thermal fluctuations via conformational transitions. Additionally, we delved into the microscopic interactions of RNA molecules, studying their sensitivity to external environments, including water, ions, and temperature. This investigation provided insights into how these factors influence RNA’s conformation and stability, particularly in relation to the specific bases present in the sequences. We believe the result of these studies will be valuable for design and engineering of low complexity RNA molecules for various biological and synthetic applications.

## METHODS and MATERIALS

### System preparation

The structures of a 20-base long single-stranded poly G, poly A, poly C and Poly U chains were generated by using X3DNA(22, 23). Each chain is placed in a cubic box of 10 nm. After that, 25 mM of ions (Na^+^ or Cl^-^) and water are added to fill the box. After preparing a solvated box containing protein, ssDNA, and ions, the system was subject to energy minimization with the conjugate-gradient algorithm using a99SBdisp-ILDN (24) and TIP3P water (25) force fields implemented in OpenMM 7.6 library (26). Following energy minimization steps, we have run 4 ns NVT and 4 ns NPT equilibration runs at temperature T = 300 K. In all these equilibration simulations, we applied a Langevin Middle integrator with a time step of 2 fs and a Nose–Hoover thermostat to maintain an average temperature with a relaxation time of 0.5 ps. We used the OpenMM’s Monte Carlo barostat to maintain an average pressure of 1 bar with a coupling time constant of 0.5 ps for NPT equilibration. All simulations used periodic boundary conditions in all directions, and the neighboring list was modified every 10 steps using a grid system with a 1 nm short range. Maintaining a NaCl concentration at 25 mg/ml allowed us to explore the impact of base chemistry on RNA conformation and dynamics (27). Experiments have shown that higher salt concentration suppresses the ability of RNA to undergo phase-transitions. Therefore we mimic experiments (28) by choosing a low salt concentration of NaCl at 25mM.

### Simulated tempering simulations

Because of RNA chains’ rugged free energy landscape, sampling conformational space of RNA often necessitates adopting enhanced sampling strategies (29–31). For sampling the conformational space of disordered biomolecules, various tempering techniques are typically employed such as Parallel tempering (PT) also known as replica exchange (REMD), Hamiltonian Replica Exchange and Simulated Tempering (ST) (32). Simulated Tempering(33–36) presents an attractive alternative allowing for simulations to be run on a single fast GPU such as V100s and A100s. Simulated Tempering enables the exploration of trajectories at different temperatures, leading to enhanced conformational sampling.

In simulated tempering, the temperature is attempted to change periodically among a ladder of discrete values, Ti (37). The probability for temperature transition from Ti to Tj follows the Metropolis criterion. weights for each temperature are required in simulated tempering and they are selected using the Wang-Landau algorithm (38). Upon successful temperature transition, the system’s momentum is scaled by √*Tj*/*Ti*. An exponentially distributed 40-rung temperature ladder is employed, with specific temperature ranges: 300K to 600K for Poly(G) and Poly(A), 300K to 520K for Poly(C), and 270K to 500K for Poly(U), at an exchange attempt of 4ps. Total production run simulation was performed over 7.5 microseconds for each chain. The average acceptance ratio over rungs of four RNAs is about Poly G : 28.3%, Poly A: 28.6%, Poly C: 43.7%, and Poly U: 32.02%.

To assess the convergence of our simulations, we employed a comprehensive approach. We initiated bootstrapping on PCA analysis (39), followed by a comparison of the landscape of the projection dPC1 and dPC2 (Figures S1 and S2). Moreover, we conducted a Cosine Content analysis for each RNA’s first five Principal Component (PC) components (36). Detailed information is provided in Supporting Information Figures S1 and S2. Collectively, these analyses demonstrate the convergence of our simulations and capturing of all major conformational transitions.

### Mapping conformational free energy landscapes of RNAs using torsional angles as features

Internal coordinates and dihedral torsional angles in particular, have been shown to be good order parameters capable of clustering flexible protein and RNA chains into structurally distinct conformations when combined with dimensionality reduction techniques such as PCA(40–42). RNA has six backbone torsional angles: *alpha, beta, gamma, delta, eps, and zeta*. Additionally, to account for the conformation of the sugar component, the glycosidic torsional angle *chi* is also considered (43). All seven of these torsional angles are illustrated in Figure 1. We apply sine and cosine transformations to each angle to transform dihedral angles from the dihedral angle to a linear metric coordinate space. In our 20-nucleotide RNA system, a total of 126 torsional angles exist. To reduce the dimensionality of our analysis, we leverage three distinct techniques: Principal Component Analysis (PCA), Time-Lagged Independent Component Analysis (TICA), and Variational Autoencoder (VAE). These techniques are applied to the sine and cosine-transformed values of all 252 dihedral angles.

**Figure 1:**
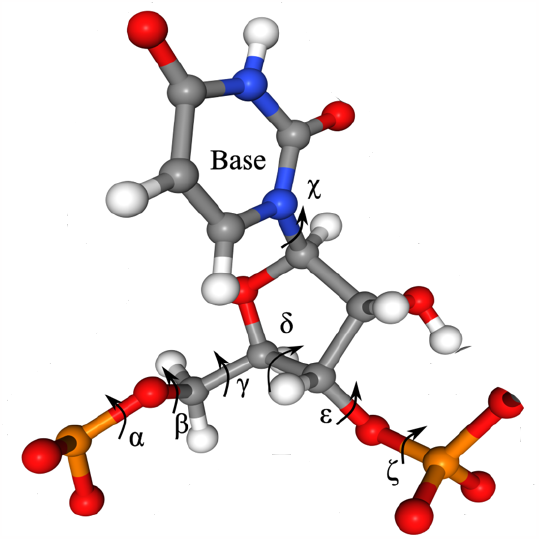
Zoomed in view of the RNA’s backbone and uracil base. We labeled dihedral angles which are used in featurization and dimensionality reduction of sampled conformational ensembles.

### Principal component analysis (PCA)

PCA is a widely used technique for reducing the dimensionality of appropriately featurized ensembles of biomolecules (44). PCA transforms the original feature space into a new set of orthogonal features called principal components (PCs) generated via linear combinations of the original features. The first principal component captures the direction of maximum variance in the data. Subsequent principal components are orthogonal to the previous ones and capture decreasing variance. PCA involves calculating the covariance matrix of the original data and then finding its eigenvalues and eigenvectors. The eigenvectors represent the directions (principal components) along which the data varies the most, and the corresponding eigenvalues represent the magnitude of variance along those directions. A key limitation of PCA is that it assumes linear relationships between correlated features and may not capture more complex nonlinear relationships.

### Time lagged independent component analysis (TICA)

TICA introduces the concept of lag-time, which determines the time span over which time-correlations are examined (45, 46). Based on a lag-time TICA identifies collective degrees of freedom that exhibit strong time-correlations. After specifying a lag-time, TICA focuses on identifying the data’s slowest and most persistent temporal patterns. TICA offers several advantages over PCA in scenarios where temporal dynamics are important. Instead of just capturing variance, TICA emphasizes capturing time-dependent structures, making it more suitable for datasets with evolving patterns. With appropriate weighting, TICA can also be applied to data generated by replica exchange and simulated tempering where natural dynamics is not preserved.

### Variational Autoencoders (VAE)

While methods like PCA and TICA are limited to linear transformations, VAEs leverage neural network architectures for capturing arbitrarily complex nonlinear relationships in the data (47). VAEs reduce the dimensionality of feature space by an encoder/decoder map. The encoder maps the input data to a lower-dimensional latent space, and the decoder reconstructs the input data from the latent representation. In VAEs, the encoder maps to mean and variance of a pre-defined distribution (usually a normal distribution). This enables VAE to generate new data points, which have the same characteristics as training data, by sampling from this distribution. The architecture of our VAE is depicted in Figure S3. It comprises three neural layers, ReLU activations applied between them. we employ the mean square loss.

## RESULTS

### Learning conformational energy landscapes of PolyA, PolyG, PolyA and PolyU RNAs

Finding an optimal dimensionality reduction method for conformationally flexible macromolecules is not a trivial task because generally the intricate inter-dependence of structural features is poorly understood. Given the conformationally flexible nature of the RNA chain without a single unique fold, internal coordinates have an advantage over cartesian coordinates. Therefore, we have featurized the conformational ensembles of RNA into 252 sine and cosine values of all torsional angles (Figure 1). Next we compare three popular dimensional reduction techniques in the space of dihedral angles, including Principal Component Analysis (PCA), Time-Lagged Independent Component Analysis (tICA), and Variational Autoencoder (VAE).

All three techniques are trained on a combined dataset of all the dihedral angles of four different RNAs sampled at different temperatures using simulated tempering. Training on a combined dataset allows machine learning techniques to (i) learn the intricate differences in conformations accompanying thermal folding/unfolding transitions of individual RNAs (ii) learn to discriminate between conformations of different RNAs at each temperature. After training on the combined dihedral angle dataset, we then projected the learned collective variables on the original data, thereby obtaining energy landscapes for different RNAs at different temperatures.

All three methods successfully delineate major conformational differences between RNAs with different bases. When it comes to differences between structurally distinct conformations of the same RNA and across temperatures, we find that VAE resolves these differences much better. In contrast, PCA and tICA generate less rugged landscapes often lumping together structurally distinct conformations (Figures 2A-C and Figures S4-S6). For instance, when considering PolyA, PCA group pseudoknot structure ‘D1’ and stem structure ‘D5’ are identical, while tICA and VAE cluster these into separate states. PCA and tICA group different knot structures ‘D3’, ‘D4’, ‘D6’, and ‘D7’ are identical, whereas VAE clusters the states into separate states within basins (Figure 3). This trend holds across the remaining bases and temperatures (Figure S4-S6). In conclusion, although PCA, tICA, and VAE succeeded in projecting landscapes for Poly G, Poly A, Poly C, and Poly U separately, VAE stands out as the superior option for delineating intricate conformational differences. This is attributed to the ability of neural networks to learn highly non-linear relationships within feature space.

**Figure 2:**
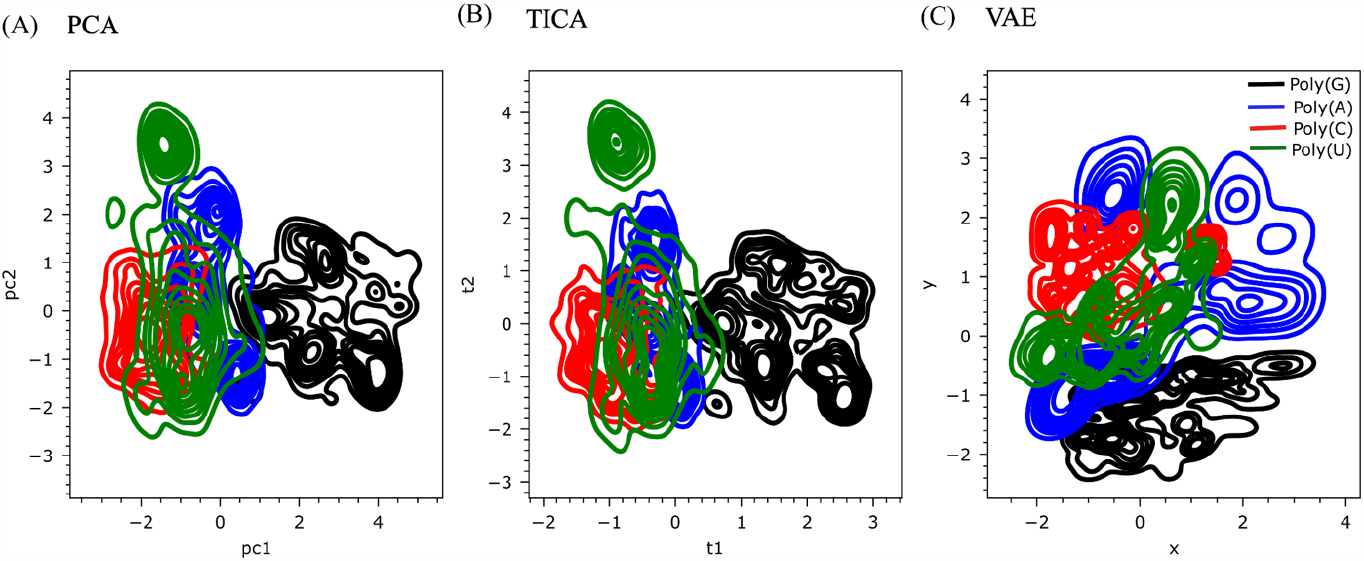
Comparative conformational energy landscapes of RNAs generated by PCA, TICA, and VAE. All three techniques are trained on a combined dataset of all the dihedral angles of four RNAs across all temperatures. Poly G is denoted in black, Poly A in blue, Poly C in red, and Poly U in green. Shows are the projections at T∼300K, from **(A)** PCA, (**B)** TICA, and **(C)** VAE.

**Figure 3:**
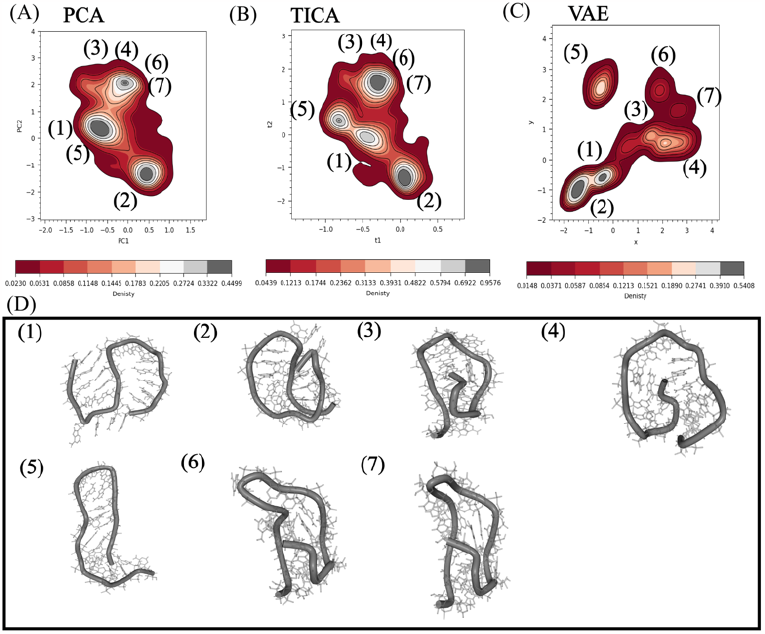
The projection of the reduced representation of dihedral angles of Poly A is presented as **(A)** PCA as a function of pc1 and pc2 **(B)** TICA as a function of t1 and t2 and **(C)** VAE as a function of x and y. **(D)** Representative structures corresponding to the different parts of the three landscapes shown on panels (A-C)

### Temperature dependence of RNA energy landscape reveals the base-specific nature of RNA unfolding

Examining temperature variation is a natural way to characterize RNA folding thermodynamics(48). Experimental techniques such as temperature-controlled optical tweezers or Isothermal Titration Calorimetry (ITC) or temperature jump experiments exploit temperature changes to investigate the thermodynamics and kinetics of RNA folding and unfolding (49–51). The atomistic details of sequence encoded conformational differences, however, remain inaccessible to most experiments. We have mapped the free energy landscape for Poly G, Poly A, Poly C, and Poly U across a range of temperatures (Figure 4 and Figure S9-S12). Through projections of the reduced components of dihedral angles, we find the impact of RNA base chemistry on the temperature dependence of the conformational dynamics of RNA chains (Figure 2A-C). When comparing different dimensionality reduction techniques in the ability to discern structurally distinct conformations, VAE emerges as a better tool for characterizing RNA’s free energy landscape. Specifically, we find multiple closely nested minima in free energy landscapes which are prominent at lower temperatures(Figures 4 and 5). Interestingly, the energy landscapes of Poly G and Poly A exhibit a slightly more intricate nature than Poly C’s, while Poly U’s landscape appears simpler than the other three (Figure 5).

**Figure 4:**
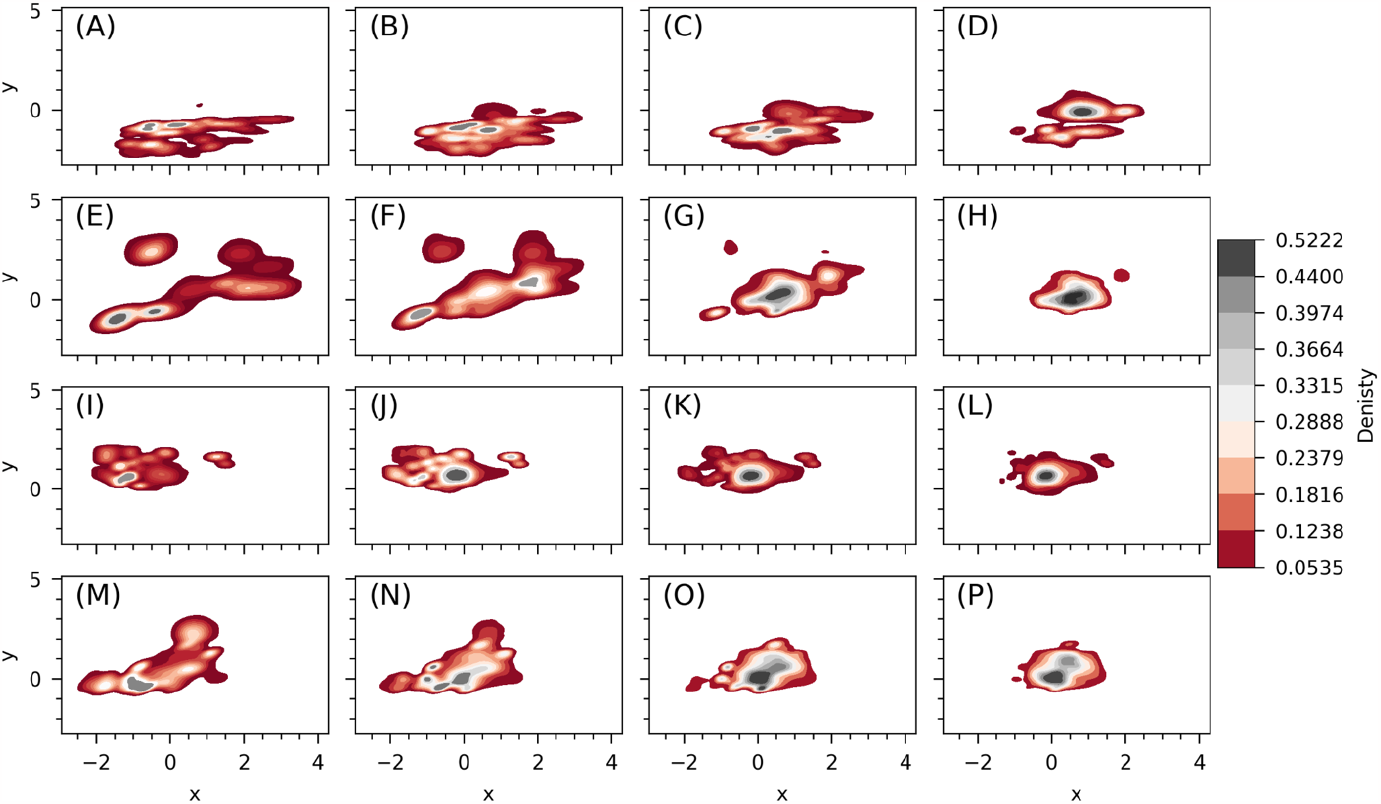
Temperature-Dependent Energy Landscapes Generated from projection of reduced components of dihedral angle from VAE on two main coordinates x and y. Panels correspond to (**A-D**) Guanine at temperatures: 300K 384K, 420K and 480K. (**E-H**) Adenine at temperatures: 300K, 352K, 384K 420K. (**I-L)** Cytosine at temperatures 300K, 326K, 350K, and 381K. (**M-P**) Uracil at temperatures: 306K, 326K, 353K and 382K.

**Figure 5:**
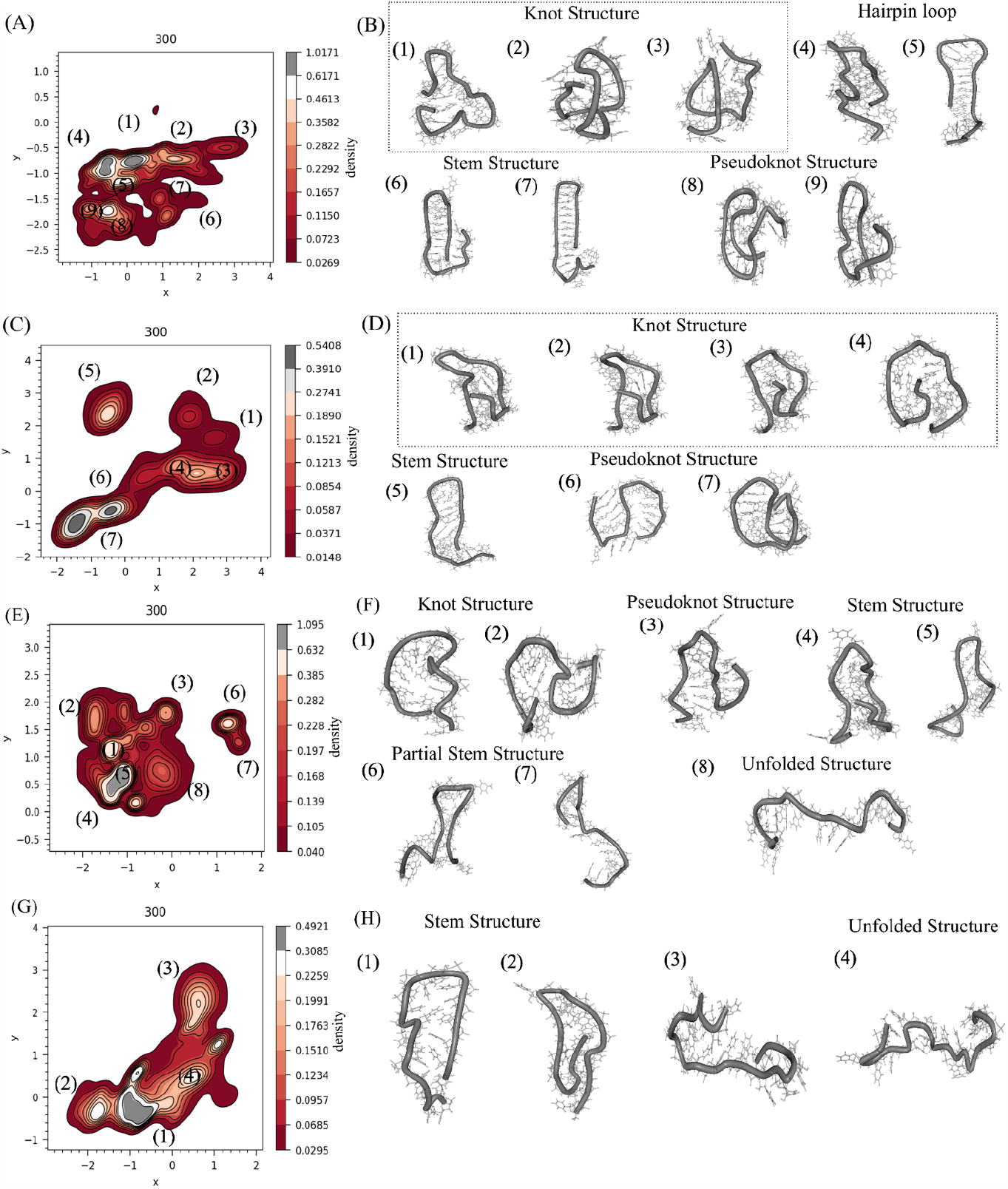
Structural diversity seen in VAE based free energy landscapes of RNA at T ∼ 300K. Panels correspond to (**A**) PolyG landscape (**B**) PolyG’s major structures (**C**) PolyA landscape (**D)** PolyA’s major structures, (**E**) PolyC landscape (**F**) PolyC’s major Structures (**G**) PolyU landscape (**H)** PolyU’s major structures.

As the temperature increases, RNA molecules access more conformations accompanied by an expanded landscape with smoother features. After crossing folding temperature, all four RNAs adopt single global minima corresponding to fully unfolded chains. Melting temperatures are dictated by base chemistry, with Poly G exhibiting thermal melting at the highest temperatures. Following Poly G are Poly A, Poly C, and Poly U. We structurally characterize the free energy landscape and reveal the role of RNA bases in shaping the type of structure they adopt becomes evident. The primary structural conformations are: (i) the unfolded state, (ii) stem structures, (iii) hairpin loops, (iv) pseudoknot structures, and (v) knot structures (52).

Interestingly, a tendency to form stem or hairpin loop structures emerges across all RNA sequences. Noticing, Poly G prefers well-ordered hairpin loops or stem structures, as illustrated in Figure 5B. Poly A leans towards forming these structures, albeit not to the extent of Poly(G). On the other hand, Poly C and Poly U display the propensity for highly mobile stem structures Where some RNA regions adopt hairpin loop or stem structures, the remaining part exhibits a mobile unfolded state (Figure 5F-H). Pseudoknot structures are also observed, with Poly A and Poly G displaying a more pronounced inclination toward this configuration, particularly in Poly A (Figure 5D). Poly G and Poly A display well-ordered structures, whereas Poly C and Poly U adopt structures featuring highly mobile segments.

The tendency to assume well-ordered structures is as follows: Poly(G) > Poly(A) > Poly(C) > Poly(U). This hierarchy underscores the varying propensities of different RNA sequences to form organized configurations. Comprehensively understanding how RNA bases shape free energy landscape necessitates more in depth exploration of microscopic interactions which include solvent and ion components.

### Temperature-dependent profiles of Hydrogen Bonding, Base Stacking, and Base Pairing

To learn the microscopic details driving base specific melting profiles and conformational landscapes of RNAs we first turn to the detailed atomistic analysis of stacking and base-pairing interactions. Before that, It’s intriguing to observe that the latent spaces generated by various unsupervised dimensionality reduction techniques exhibit similarities between Poly G and Poly C, as well as between Poly A and Poly U. It appears that Poly C and Poly U are milder counterparts of Poly G and Poly A, respectively. Therefore, we have looked into the interactions between (i) Bases, (ii) Base and phosphate group, and (iii) Base and sugars by plotting the radial distribution function for all four RNA chains: Poly G, Poly A, Poly C, and PolyU.

We find that Poly G and Poly C exhibit similar behaviors in the interaction between base and phosphate group at two specific distances, approximately 4 Angstroms and 7 Angstroms, with Poly G showing slightly more extensive interactions (Figure S15A). Conversely, Poly A and Poly U lack a pronounced peak at around 4 Angstroms in this interaction (Figure S15B). Likewise, in the interaction between bases and sugars, Poly G and Poly C display similar behaviors, while Poly A and Poly U exhibit more pronounced interactions. Poly U displays sharp peaks at nearly 4 Angstroms (Figure S15C and S15D).In the context of base-base interactions, as expected, Poly G and Poly A exhibit stronger interactions than Poly U and Poly C (Figure S16). The absence of strong base-base interaction in Poly C and Poly U characterizes them as milder counterparts of Poly G and Poly A.

We use radius of gyration (Rg) against temperature as a measure of RNA folding/unfolding transitions (Figure 6a). Radius of gyration displays a sigmoidal shape indicative of two-state thermodynamics of folding. Consistent with temperature dependence of free energy landscapes, melting temperatures follow Poly(G) > Poly(A) > Poly(C) > Poly(U) order. We break down the intramolecular RNA contacts into hydrogen bonding and base stacking groups. Hydrogen bonding includes contacts between base pairs, between phosphate and base, as well as between sugar and base. Stacking includes intra RNA stacking interactions. We computed the average number of hydrogen bonds against temperature (Figure 6B), the average number of stacking interactions against temperature (Figure 6C), and the average number of base pairs against temperature (Figure 6D). While all three plots exhibit folding transitions with a sigmoid shape, they do not mirror each other precisely. This discrepancy underscores the pivotal role of specific interactions within each system.

**Figure 6:**
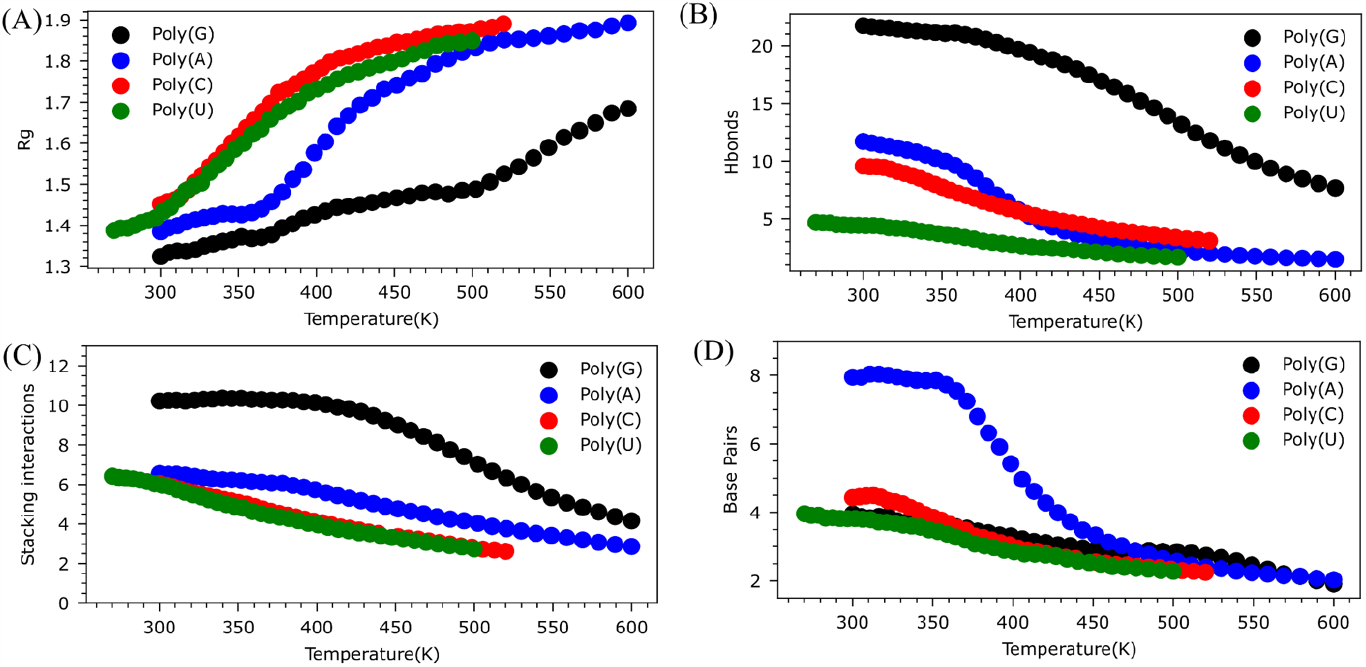
Quantifying microscopic drivers of RNA (un)folding for all four RNAs across temperatures. Guanine is denoted in black, adenine in blue, cytosine in red, and uracil in green. (**A**) Average Rg (nm) vs. temperature (**B**) Average number of hydrogen bonds vs. temperature, (**C**) Stacking interaction vs. temperature, and (**D**) Average number of base pairs vs. temperature.

Poly(G) notably displays the highest count of hydrogen bonds and stacking interactions. Purines, such as Guanine and Adenine, can clearly engage in more stacking interactions compared to pyrimidines, Cytosine and Uracil (Figure 6C). Number of hydrogen bonds follows: Poly(G) > Poly(A) > Poly(C) > Poly(U) order (Figure 6B and Figure S17) while the stacking interactions follows: Poly(G) > Poly(A) > Poly(C) = Poly(U) order. Interestingly, Poly(A) can readily form base pairs (Figure 6D), a capability not as occurred in the other bases. These microscopic interactions contribute to the formation of a range of structural motifs, ultimately influencing the ruggedness of the free energy landscape (Figure 4-5) Although the average interaction counts provide insight into why the free energy landscape is base-dependent, to quantify influence of different interactions we compute Pearson correlation between Rg and non-bonded interaction types; hydrogen bonds and stacking interactions across temperatures.

Poly(G) adopts well-organized stem or hairpin loop structures as a result of strong stacking interactions among its bases, supported by the formation of hydrogen bonds between the NH2 and NH groups of the base and the phosphate groups (Figure 7C). The correlation between Rg and stacking sharply decreases with increasing temperature, while the correlation between Rg and hydrogen bonds remains relatively unaffected (Figure 7 A & B). This observation implies that stacking interactions primarily govern the Poly(G) dynamics rather than hydrogen bonds.

**Figure 7:**
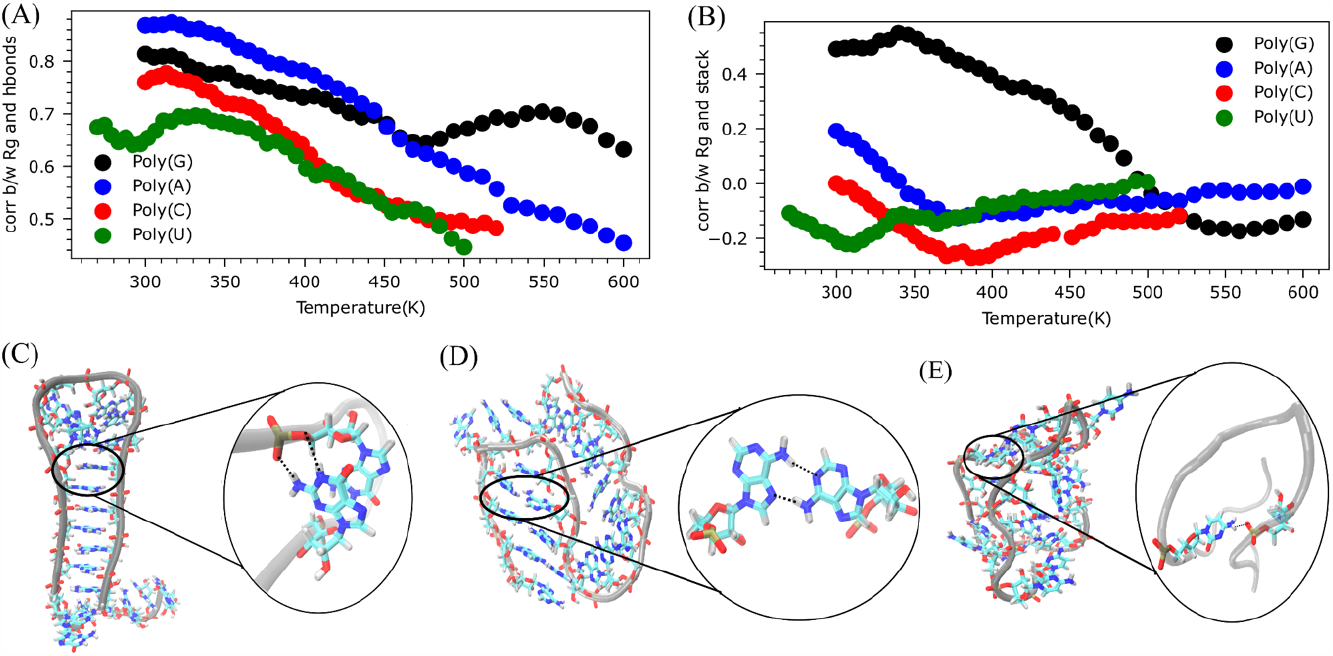
Correlation of hydrogen bonding patterns with conformational dynamics of RNAs (**A**) Pearson Correlation Between Rg and Hydrogen Bond Count in RNA across temperatures for all four RNAs. (**B**) Correlation Between Rg and Stacking Interactions in RNA across temperatures. (**C**) Illustration of the hairpin loop structure in Poly G, highlighting stacking interactions between bases and hydrogen bonds between bases and phosphate Backbone on the right side. (**D**) The pseudoknot structure in Poly A is represented with base pairing shown on the right side. (**E**) Depiction of the knot structure in Poly C, emphasizing the hydrogen bond between the base and phosphate backbone on the right side.

Poly(A)’s ability to form base pairs (Figure 7C) leads to numerous base-paired stems, hairpin loops, and pseudoknot structures. This is distinct from Poly(A), where well-ordered stacking interactions are predominant. Figure 7C highlights that Poly(A) tends to form pseudoknot structures, attributed to base-paired stems and NH_2_ groups of parallel loop bases forming hydrogen bonds with stem phosphate groups. The correlation between Rg and hydrogen bonds sharply increases with temperature, while the correlation between Rg and stacking interactions remains relatively stable (Figure 7A & B). This outcome emphasizes that hydrogen bonds primarily dictate the dynamics of Poly(A) rather than stacking interactions. In Poly(C), hydrogen bonds between the NH_2_ of the base and phosphate groups emerge as the predominant interactions (Figure 7E, forming flexible, mobile stems, hairpin loops, or pseudoknot structures. Conversely, in Poly(U), the absence of dominant interactions contributes to the highly mobile nature of its structure compared to the other three bases.

In sum from atomistic analysis of non-bonded contacts it is clear that chemical structures of the nitrogen base plays a significant role in shaping its conformational preferences. This influence is exerted through interactions of the nitrogen base with itself, the phosphate group, and the ribose sugar.

### Impact of temperature-dependent hydration and ionic condensation on RNA conformational dynamics

The conformationally extended RNA forms form an extensive network of hydrogen bonds and due to high charge density, are screened significantly by the ionic environment. To discern the base specific nature of RNA conformational preferences, it is necessary to dissect the role of ion and water-mediated interactions. We quantify the number of water molecules within 4 Angstroms of RNA at various temperatures. As anticipated, with increasing temperature, RNA unfolds, coinciding with a rise in the average count of water molecules surrounding it (Figure 8A-C). Furthermore, we calculate correlations between Radius of Gyration and the number of water molecules surrounding RNA at different temperatures, as well as correlations between the number of water molecules and the number of hydrogen bonds at various temperatures.

**Figure 8:**
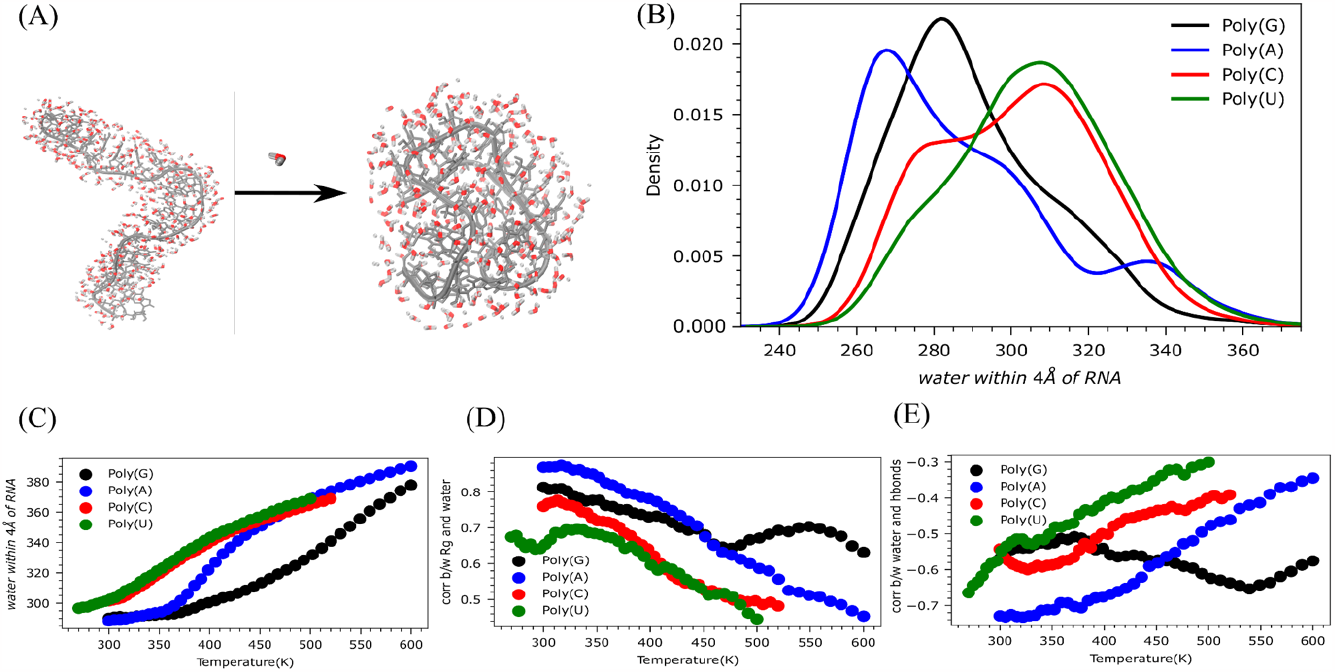
Quantifying the role of water on RNA conformational dynamics (**A**) Depiction of water expulsion as RNA folds into a knot structure. (**B**) Distribution of the amount of water within a 4 Angstrom of RNA for all four RNAs at T∼ 300K (**C**) Average amount of water within 4 Angstrom across temperatures for all four RNAs. (**D**) Pearson Correlation Between Rg and number of water molecules within 4 Angstroms of RNA across temperatures for all four RNAs. (**E**) Correlation Between the number of water molecules within 4 Angstroms of RNA and the number of hydrogen bonds in RNA across temperatures for all four RNAs.

The correlation between Rg and the number of water molecules around RNA proves to be highly positive (as depicted in Figure 8D). This positive correlation points to entropy gain resulting from water expulsion driven RNA folding (Figure 8A).

The number of water molecules around RNA at T=300K is as follows: Poly(A) < Poly(G) < Poly(C) < Poly(U) (Figure 8B), whereas Rg at T=300K is: Poly(G) < Poly(A) < Poly(C) < Poly(U) (Figure S13). This suggests that while Poly(G) has the lowest Rg, Poly(A) possesses the lowest count of water molecules around it. Additionally, correlation between Rg and the average number of water molecules is: Poly(A) > Poly(G) > Poly(C) > Poly(U) (Figure 8D), and correlation between water molecules around RNA and the number of hydrogen bonds is: Poly(A) << Poly(G) = Poly(U) = Poly(C) (Figure 8E).

These observations indicate that Poly(A) needs to repel a substantial number of water molecules in order to adopt folded conformations. Conversely, Poly(G) folds with less water repulsion, implying that the folding of Poly(G) is primarily driven by enthalpy as opposed to Poly(A). These correlations remain stable with increasing temperature in contrast to the other three, underscoring that water’s influence on Poly(G) is comparatively limited due to its dynamics being driven by stacking interactions.

For the other three sequences, Poly(A), Poly(U), and Poly(C), these two correlations exhibit a steep decline with rising temperature (Figure 8D-E), emphasizing the significant influence of water on their conformational behavior. Finally, when examining the ionic environment around RNA chains, we find a negative correlation between Rg and the number of ions, implying that as RNA folds, an increasing number of cations gather around to counteract the repulsive force generated by the negatively charged backbone of RNA (Figure S14 A-C).

Simultaneously, as the temperature rises, the number of ions around RNA increases due to greater exposure (Figure S14 B).

## Discussion

The conformational dynamics of RNA and its affinity to engage in multivalent interactions underlies the ability of RNA to form biomolecular condensates(20, 21, 28, 53–55). Unlocking the relationship between RNA’s folding free energy landscapes and affinity for biomolecular condensate formation is therefore essential for understanding the sequence-dynamics-function relationships of RNA. In this work, we have employed simulated tempering and deep generative learning to map the energy landscapes of homopolymeric RNAs across a range of temperatures encompassing fully folded and unfolded conformational states. We have dissected the key microscopic interactions contributing to distinct folding energy landscapes of RNAs with different base makeups. Using unsupervised learning techniques in the space of backbone torsion angles of RNA we mapped conformational energy landscapes for different temperatures which encompasses folded and unfolded states. We found Variational Autoencoders superior in their ability to discriminate structurally distinct conformations of RNA. Analysis of the energy landscape of Poly G, Poly A, Poly C and Poly U at different temperatures reveals numerous insights on the nature of base chemistry on conformational dynamics of RNA. We are able to dissect behavior common to polyphosphate groups from base specific chemistry. We find that RNA tends to form stems, hairpin loops, or pseudoknot structures regardless of the base. Simulations predicate the thermal melting trends as follows: Poly G > Poly A> Poly C > PolyU.

Unsupervised learning techniques are able to effectively differentiate between structurally heterogeneous conformational ensemble RNAs with different bases. We find that stacking interactions primarily drive the dynamics of Poly(G). As a result Poly(G) assumes well-organized stem or hairpin loop structures, a consequence of robust stacking interactions among its bases, further strengthened by the formation of hydrogen bonds between bases and the backbone (11, 56). This type of stem formation in RNA is novel in our current understanding. PolyA dynamics, on the other hand, is driven by hydrogen bonds, and its ability to form base pairs has been documented in the past(11). This leads adenine to form a stem and pseudoknot structure. In PolyC, hydrogen bonds between the NH2 of the base and phosphate groups emerge as the predominant interactions, allowing for the formation of flexible, mobile stems, hairpin loops, or pseudoknot structures. In contrast, in PolyU, the absence of dominant interactions contributes to the highly mobile nature of its structure compared to the other three bases. The inability of Poly(U) to form cross-links aligns with recent experiments from Banerjee group, illustrating that multivalent ions like spermine or Mg^2+^ are essential for Poly(U) to undergo phase separation (17, 18). Mg^2+^ ions facilitate intermolecular connections through chelation, emerging as a dominant interaction driving phase separation.

A recent study by Jain revealed that RNA’s propensity to undergo multivalent base-pairing can induce gelation without proteins. This phenomenon was observed in CAG, CUG, and GGGGCC repeats (20, 57, 58). Nonetheless, the phase separation by Poly(G), Poly(A), Poly(C), and Poly(U) shows that sequence-specific base pairing isn’t necessary for phase separation (18). RNA folding is inevitably accompanied by the expulsion of hydration layers from RNA (59). We find distinct signatures of water hydrogen bonding, which contributes to the folding profiles of RNA. Hydrogen bond determines the dynamics of poly A, poly U, and poly C, leading water to influence its dynamics significantly. In contrast, water dynamics influence less Poly G dynamics as stacking interactions drive them. Interestingly, poly A has to repel a substantial amount of water to fold compared to Poly G, showing that poly G is enthalpically driven as compared to Poly A.

To sum up, our study of folding energy landscapes of homopolymeric RNA provide fresh insights into RNA behavior, encompassing RNA folding, phase separation, and other related phenomena. We establish that experimentally observed temperature-driven shifts in metastable state populations align well with the experimental phase diagrams for homopolymeric RNAs. The work establishes a microscopic framework to reason about base specific RNA propensity for phase-separation. We believe our work will be valuable for the design of novel RNA sensors for biological and synthetic applications.

## DATA AVAILABILITY

All data is available in the manuscript and supplementary information file.

## SUPPLEMENTARY DATA

Supplementary Data are available at NAR online.

## AUTHOR CONTRIBUTIONS

Vysakh Ramachandran: Conceptualization, Analysis, Methodology, Writing. Davit Potoyan: Conceptualization, Writing.

## ACKNOWLEDGEMENTS

We thank Prof Priya Banerjee and Dr Ibraheem Alshareedah for fruitful discussions.

## REFERENCES

1. Sharp, S.J., Schaack, J., Cooley, L., Burke, D.J. and Soil, D. (1985) Structure and Transcription of Eukaryotic tRNA Gene. Crit. Rev. Biochem. Mol. Biol., 19, 107–144.

2. Kruger, K., Grabowski, P.J., Zaug, A.J., Sands, J., Gottschling, D.E. and Cech, T.R. (1982) Self-splicing RNA: autoexcision and autocyclization of the ribosomal RNA intervening sequence of Tetrahymena. Cell, 31, 147–157.

3. Emilsson, G.M., Nakamura, S., Roth, A. and Breaker, R.R. (2003) Ribozyme speed limits. RNA, 9, 907–918.

4. Carter, A.P., Clemons, W.M., Brodersen, D.E., Morgan-Warren, R.J., Wimberly, B.T. and Ramakrishnan, V. (2000) Functional insights from the structure of the 30S ribosomal subunit and its interactions with antibiotics. Nature, 407, 340–348.

5. Walter, N.G. and Maquat, L.E. (2018) Introduction-RNA: From Single Molecules to Medicine. Chem. Rev., 118, 4117–4119.

6. Kührová, P., Mlýnský, V., Otyepka, M., Šponer, J. and Banáš, P. (2023) Sensitivity of the RNA structure to ion conditions as probed by molecular dynamics simulations of common canonical RNA duplexes. J. Chem. Inf. Model., 63, 2133–2146.

7. Ramachandran, V., Mainan, A. and Roy, S. (2022) Dynamic effects of the spine of hydrated magnesium on viral RNA pseudoknot structure. Phys. Chem. Chem. Phys., 24, 24570–24581.

8. Fischer, N.M., Polêto, M.D., Steuer, J. and van der Spoel, D. (2018) Influence of Na+ and Mg2+ ions on RNA structures studied with molecular dynamics simulations. Nucleic Acids Res., 46, 4872–4882.

9. Mathez, G. and Cagno, V. (2023) Small Molecules Targeting Viral RNA. Int. J. Mol. Sci., 24.

10. Childs-Disney, J.L., Yang, X., Gibaut, Q.M.R., Tong, Y., Batey, R.T. and Disney, M.D. (2022) Targeting RNA structures with small molecules. Nat. Rev. Drug Discov., 21, 736–762.

11. Zarudnaya, M.I., Kolomiets, I.M., Potyahaylo, A.L. and Hovorun, D.M. (2019) Structural transitions in poly(A), poly(C), poly(U), and poly(G) and their possible biological roles. J. Biomol. Struct. Dyn., 37, 2837–2866.

12. Napthine, S., Treffers, E.E., Bell, S., Goodfellow, I., Fang, Y., Firth, A.E., Snijder, E.J. and Brierley, I. (2016) A novel role for poly(C) binding proteins in programmed ribosomal frameshifting. Nucleic Acids Res., 44, 5491–5503.

13. Kolykhalov, A.A., Feinstone, S.M. and Rice, C.M. (1996) Identification of a highly conserved sequence element at the 3’ terminus of hepatitis C virus genome RNA. J. Virol., 70, 3363–3371.

14. Long, Q., Hua, Y., He, L., Zhang, C., Sui, S., Li, Y., Qiu, H., Tian, T., An, X., Luo, G., et al. (2020) Poly(U) binding splicing factor 60 promotes renal cell carcinoma growth by transcriptionally upregulating telomerase reverse transcriptase. Int. J. Biol. Sci., 16, 3002–3017.

15. Passmore, L.A. and Coller, J. (2022) Roles of mRNA poly(A) tails in regulation of eukaryotic gene expression. Nat. Rev. Mol. Cell Biol., 23, 93–106.

16. Brouze, A., Krawczyk, P.S., Dziembowski, A. and Mroczek, S. (2023) Measuring the tail: Methods for poly(A) tail profiling. Wiley Interdiscip. Rev. RNA, 14, e1737.

17. Pullara, P., Alshareedah, I. and Banerjee, P.R. (2022) Temperature-dependent reentrant phase transition of RNA-polycation mixtures. Soft Matter, 18, 1342–1349.

18. Wadsworth, G.M., Zahurancik, W.J., Zeng, X., Pullara, P., Lai, L.B., Sidharthan, V., Pappu, R.V., Gopalan, V. and Banerjee, P.R. (2022) RNAs undergo phase transitions with lower critical solution temperatures. bioRxiv, 10.1101/2022.10.17.512593.

19. Malhotra, I. and Potoyan, D.A. (2023) Re-entrant transitions of locally stiff RNA chains in the presence of polycations leads to gelated architectures. Soft Matter, 19, 5622–5629.

20. Nguyen, H.T., Hori, N. and Thirumalai, D. (2022) Condensates in RNA repeat sequences are heterogeneously organized and exhibit reptation dynamics. Nat. Chem., 14, 775–785.

21. Maity, H., Nguyen, H.T., Hori, N. and Thirumalai, D. (2023) Odd–even disparity in the population of slipped hairpins in RNA repeat sequences with implications for phase separation. Proceedings of the National Academy of Sciences, 120, e2301409120.

22. Zheng, G., Lu, X.-J. and Olson, W.K. (2009) Web 3DNA--a web server for the analysis, reconstruction, and visualization of three-dimensional nucleic-acid structures. Nucleic Acids Res., 37, W240–6.

23. Lu, X.-J. and Olson, W.K. (2003) 3DNA: a software package for the analysis, rebuilding and visualization of three-dimensional nucleic acid structures. Nucleic Acids Res., 31, 5108–5121.

24. Robustelli, P., Piana, S. and Shaw, D.E. (2018) Developing a molecular dynamics force field for both folded and disordered protein states. Proc. Natl. Acad. Sci. U. S. A., 115, E4758–E4766.

25. Jorgensen, W.L., Chandrasekhar, J., Madura, J.D., Impey, R.W. and Klein, M.L. (1983) Comparison of simple potential functions for simulating liquid water. J. Chem. Phys., 79, 926–935.

26. Eastman, P., Swails, J., Chodera, J.D., McGibbon, R.T., Zhao, Y., Beauchamp, K.A., Wang, L.-P., Simmonett, A.C., Harrigan, M.P., Stern, C.D., et al. (2017) OpenMM 7: Rapid development of high performance algorithms for molecular dynamics. PLoS Comput. Biol., 13, e1005659.

27. Plumridge, A., Andresen, K. and Pollack, L. (2020) Visualizing Disordered Single-Stranded RNA: Connecting Sequence, Structure, and Electrostatics. J. Am. Chem. Soc., 142, 109–119.

28. Alshareedah, I., Kaur, T., Ngo, J., Seppala, H., Kounatse, L.-A.D., Wang, W., Moosa, M.M. and Banerjee, P.R. (2019) Interplay between Short-Range Attraction and Long-Range Repulsion Controls Reentrant Liquid Condensation of Ribonucleoprotein–RNA Complexes. J. Am. Chem. Soc., 141, 14593–14602.

29. Chakraborty, D., Collepardo-Guevara, R. and Wales, D.J. (2014) Energy landscapes, folding mechanisms, and kinetics of RNA tetraloop hairpins. J. Am. Chem. Soc., 136, 18052–18061.

30. Bottaro, S., Banáš, P., Šponer, J. and Bussi, G. (2016) Free Energy Landscape of GAGA and UUCG RNA Tetraloops. J. Phys. Chem. Lett., 7, 4032–4038.

31. Zerze, G.H., Piaggi, P.M. and Debenedetti, P.G. (2021) A Computational Study of RNA Tetraloop Thermodynamics, Including Misfolded States. J. Phys. Chem. B, 125, 13685–13695.

32. Qi, R., Wei, G., Ma, B. and Nussinov, R. (2018) Replica Exchange Molecular Dynamics: A Practical Application Protocol with Solutions to Common Problems and a Peptide Aggregation and Self-Assembly Example. In Nilsson, B.L., Doran, T.M. (eds), Peptide Self-Assembly: Methods and Protocols. Springer New York, New York, NY, pp. 101–119.

33. Sousa, C.F., Becker, R.A., Lehr, C.-M., Kalinina, O.V. and Hub, J.S. (2023) Simulated Tempering-Enhanced Umbrella Sampling Improves Convergence of Free Energy Calculations of Drug Membrane Permeation. J. Chem. Theory Comput., 19, 1898–1907.

34. Park, S. and Pande, V.S. (2007) Choosing weights for simulated tempering. Phys. Rev. E Stat. Nonlin. Soft Matter Phys., 76, 016703.

35. Nagai, T. and Okamoto, Y. (2012) Simulated tempering and magnetizing: Application of two-dimensional simulated tempering to the two-dimensional Ising model and its crossover. Phys. Rev. E, 86, 056705.

36. Pan, A.C., Weinreich, T.M., Piana, S. and Shaw, D.E. (2016) Demonstrating an Order-of-Magnitude Sampling Enhancement in Molecular Dynamics Simulations of Complex Protein Systems. J. Chem. Theory Comput., 12, 1360–1367.

37. Chodera, J.D. and Shirts, M.R. (2011) Replica exchange and expanded ensemble simulations as Gibbs sampling: simple improvements for enhanced mixing. J. Chem. Phys., 135, 194110.

38. Wang, F. and Landau, D.P. (2001) Efficient, multiple-range random walk algorithm to calculate the density of states. Phys. Rev. Lett., 86, 2050–2053.

39. Romo, T.D. and Grossfield, A. (2011) Block Covariance Overlap Method and Convergence in Molecular Dynamics Simulation. J. Chem. Theory Comput., 7, 2464–2472.

40. Sittel, F. and Stock, G. (2018) Perspective: Identification of collective variables and metastable states of protein dynamics. J. Chem. Phys., 149, 150901.

41. Riccardi, L., Nguyen, P.H. and Stock, G. (2009) Free-energy landscape of RNA hairpins constructed via dihedral angle principal component analysis. J. Phys. Chem. B, 113, 16660–16668.

42. Miner, J.C., Chen, A.A. and García, A.E. (2016) Free-energy landscape of a hyperstable RNA tetraloop. Proc. Natl. Acad. Sci. U. S. A., 113, 6665–6670.

43. Richardson, J.S., Schneider, B., Murray, L.W., Kapral, G.J., Immormino, R.M., Headd, J.J., Richardson, D.C., Ham, D., Hershkovits, E., Williams, L.D., et al. (2008) RNA backbone: consensus all-angle conformers and modular string nomenclature (an RNA Ontology Consortium contribution). RNA, 14, 465–481.

44. Glielmo, A., Husic, B.E., Rodriguez, A., Clementi, C., Noé, F. and Laio, A. (2021) Unsupervised Learning Methods for Molecular Simulation Data. Chem. Rev., 121, 9722–9758.

45. Naritomi, Y. and Fuchigami, S. (2011) Slow dynamics in protein fluctuations revealed by time-structure based independent component analysis: the case of domain motions. J. Chem. Phys., 134, 065101.

46. Molgedey, L. and Schuster, H.G. (1994) Separation of a mixture of independent signals using time delayed correlations. Phys. Rev. Lett., 72, 3634–3637.

47. Pinheiro Cinelli, L., Araújo Marins, M., Barros da Silva, E.A. and Lima Netto, S. (2021) Variational Autoencoder. In Cinelli, L.P., Marins, M.A., Barros da Silva, E.A., Netto, S.L. (eds), Variational Methods for Machine Learning with Applications to Deep Networks. Springer International Publishing, Cham, pp. 111–149.

48. Ma, H., Proctor, D.J., Kierzek, E., Kierzek, R., Bevilacqua, P.C. and Gruebele, M. (2006) Exploring the energy landscape of a small RNA hairpin. J. Am. Chem. Soc., 128, 1523–1530.

49. Suurkuusk, J., Alvarez, J., Freire, E. and Biltonen, R. (1977) Calorimetric determination of the heat capacity changes associated with the conformational transitions of polyriboadenylic acid and polyribouridylic acid. Biopolymers, 16, 2641–2652.

50. Feig, A.L. (2007) Applications of isothermal titration calorimetry in RNA biochemistry and biophysics. Biopolymers, 87, 293–301.

51. Mikulecky, P.J. and Feig, A.L. (2004) Heat capacity changes in RNA folding: application of perturbation theory to hammerhead ribozyme cold denaturation. Nucleic Acids Res., 32, 3967–3976.

52. Šponer, J., Bussi, G., Krepl, M., Banáš, P., Bottaro, S., Cunha, R.A., Gil-Ley, A., Pinamonti, G., Poblete, S., Jurečka, P., et al. (2018) RNA Structural Dynamics As Captured by Molecular Simulations: A Comprehensive Overview. Chem. Rev., 118, 4177–4338.

53. Roden, C. and Gladfelter, A.S. (2020) RNA contributions to the form and function of biomolecular condensates. Nat. Rev. Mol. Cell Biol., 10.1038/s41580-020-0264-6.

54. Kimchi, O., King, E.M. and Brenner, M.P. (2023) Uncovering the mechanism for aggregation in repeat expanded RNA reveals a reentrant transition. Nat. Commun., 14, 1–9.

55. Wiedner, H.J. and Giudice, J. (2021) It’s not just a phase: function and characteristics of RNA-binding proteins in phase separation. Nat. Struct. Mol. Biol., 28, 465–473.

56. Rice, J., Lafleur, L., Medeiros, G.C. and Thomas, G.J., Jr (1973) Raman studies of nucleic acids. IX: A salt-induced structural transition in poly(rG). J. Raman Spectrosc., 1, 207–215.

57. Jain, A. and Vale, R.D. (2017) RNA phase transitions in repeat expansion disorders. Nature, 546, 243–247.

58. Ma, Y., Li, H., Gong, Z., Yang, S., Wang, P. and Tang, C. (2022) Nucleobase Clustering Contributes to the Formation and Hollowing of Repeat-Expansion RNA Condensate. J. Am. Chem. Soc., 144, 4716–4720.

59. Templeton, C. and Elber, R. (2018) Why Does RNA Collapse? The Importance of Water in a Simulation Study of Helix-Junction-Helix Systems. J. Am. Chem. Soc., 140, 16948–16951.

