## Supporting Information for "Mapping energy landscapes of homopolymeric RNAs via simulated tempering and deep unsupervised learning"

**Measuring convergence to equilibrium distributions:**

Bootstrapping is a statistical technique used to measure statistical error in data analysis. It involves the random selection of two separate sets of data points, followed by a comparison between them. We randomly selected two trajectories comprising 10,000 frames from the entire trajectory dataset. Subsequently, independent dPCA (dihedral Principal component analysis) analyses were conducted on these randomly sampled trajectories. A comparison between the projection landscapes of PC1 and PC2 obtained from these two distinct datasets followed for all RNAs at T~300K (Figure S1 and S2).

Additionally, we calculated Covariance Overlap and PCA Similarity Factor Between these two. Remarkably, we observed a substantial degree of overlap between these two datasets, indicating the robustness and consistency of our analysis.

In addition, we conducted a Cosine Content analysis on dPCA for all four RNA molecules. The Cosine Content of each RNA's first five PC components is shown in the table below. These values range from 0 (indicating no similarity to a cosine shape) to 1 (representing a perfect cosine shape). If values below 0.7 do not necessarily indicate inadequate sampling. Importantly, our simulations exhibited convergence.

#### Cosine Content analysis:

**Guanine**
Cosine content for PC 1 = 0.422
Cosine content for PC 2 = 0.257
Cosine content for PC 3 = 0.056
Cosine content for PC 4 = 0.251
Cosine content for PC 5 = 0.208

**Adenine**
Cosine content for PC 1 = 0.143
Cosine content for PC 2 = 0.733
Cosine content for PC 3 = 0.233
Cosine content for PC 4 = 0.139
Cosine content for PC 5 = 0.207

**Cytosine**
Cosine content for PC 1 = 0.129
Cosine content for PC 2 = 0.001
Cosine content for PC 3 = 0.172
Cosine content for PC 4 = 0.038
Cosine content for PC 5 = 0.001

**Uracil**
Cosine content for PC 1 = 0.355
Cosine content for PC 2 = 0.013
Cosine content for PC 3 = 0.056
Cosine content for PC 4 = 0.002
Cosine content for PC 5 = 0.099

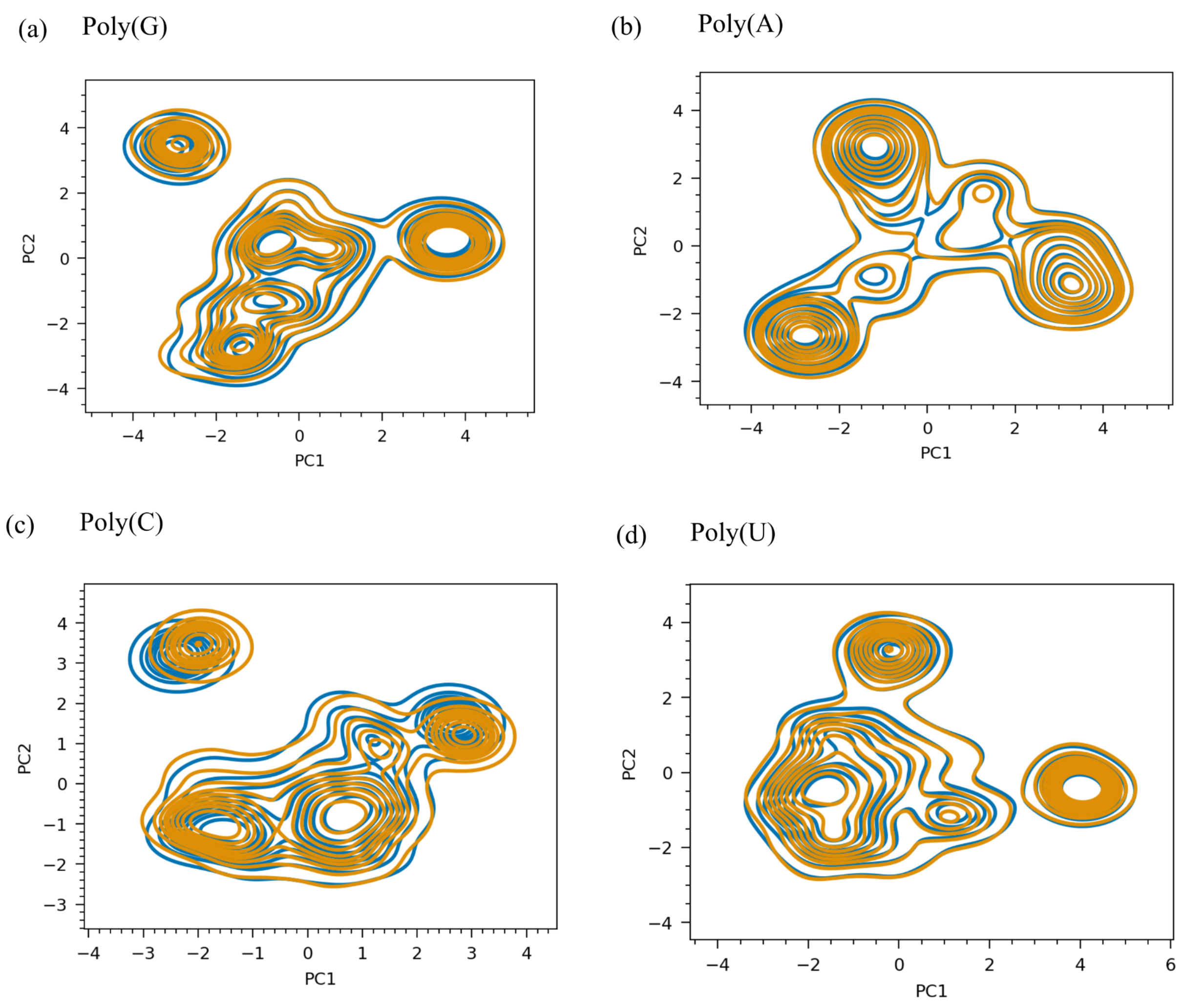


**Figure S1**: Comparison of landscape of the projection dPC1 and dPC2 from Two Bootstrapped Trajectories, Each Comprising 10,000 Frames, for (a) Guanine, (b) Adenine, (c) Cytosine, and (d) Uracil at T~300K.

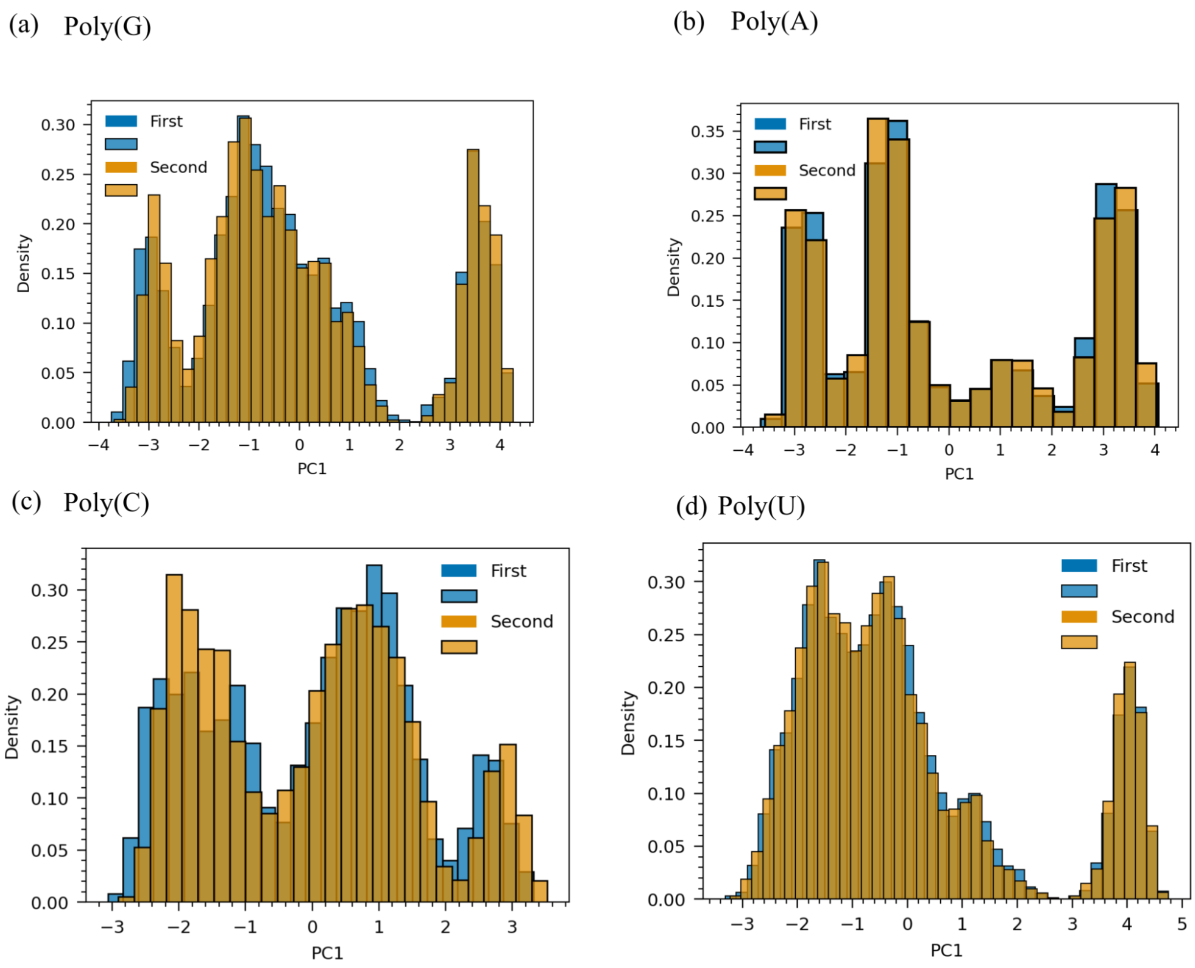


**Figure S2**: Figure S1: Comparison of dPC1 from Two Bootstrapped Trajectories, Each Comprising 10,000 Frames, for (a) Poly(G), (b) Poly(A), (c) Poly(C), and (d) Poly(U) at T~300K.

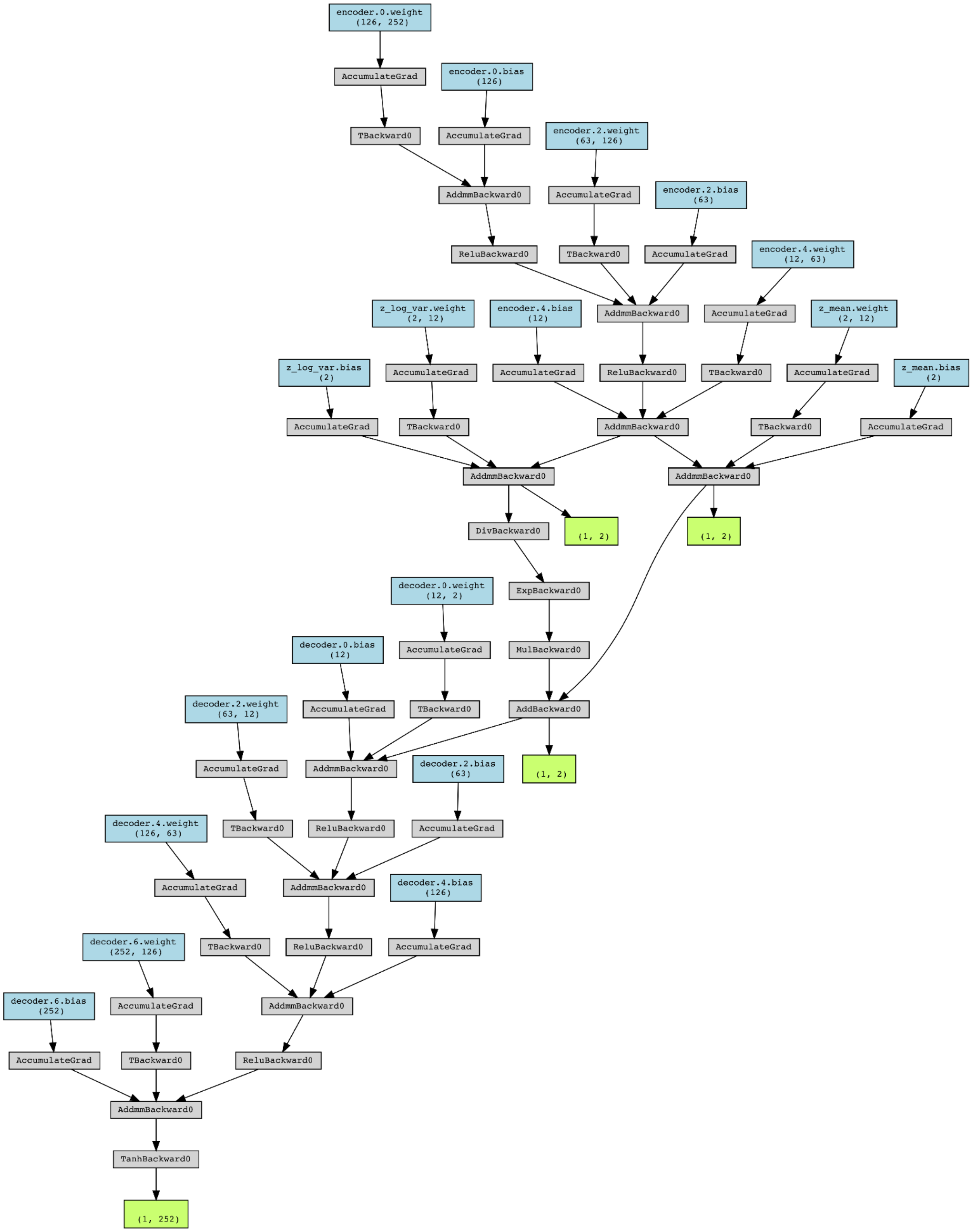


**Figure S3**: Architecture of the Variational Autoencoder (VAE).

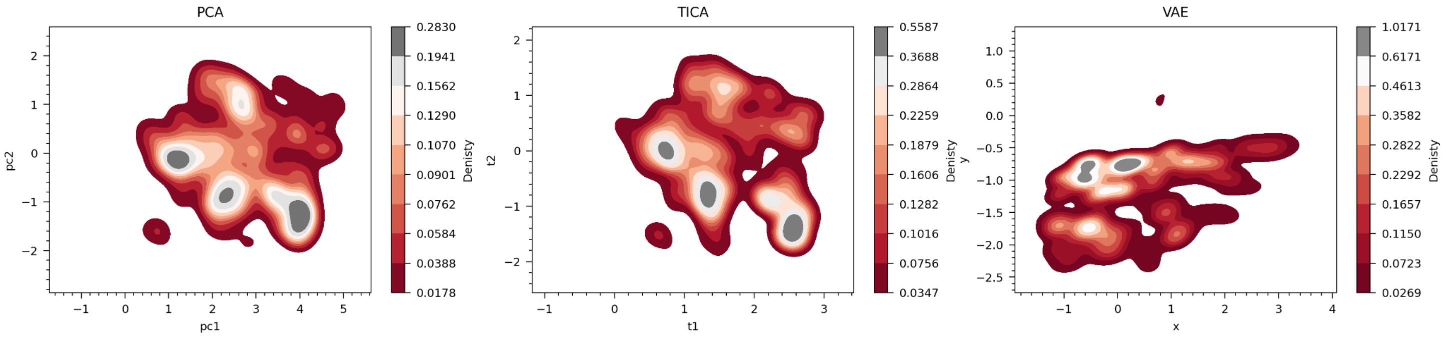


**Figure S4:** The projection of reduced components of dihedral angles of Poly(G) is presented as (a) PCA, (b) TICA, and (c) VAE.

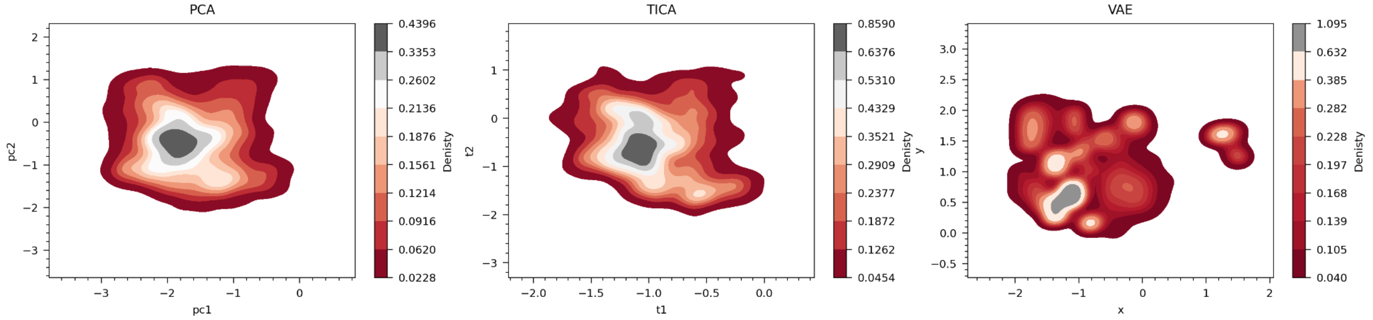


**Figure S5:** The projection of reduced components of dihedral angles of Poly(C) is presented as (a) PCA, (b) TICA, and (c) VAE.

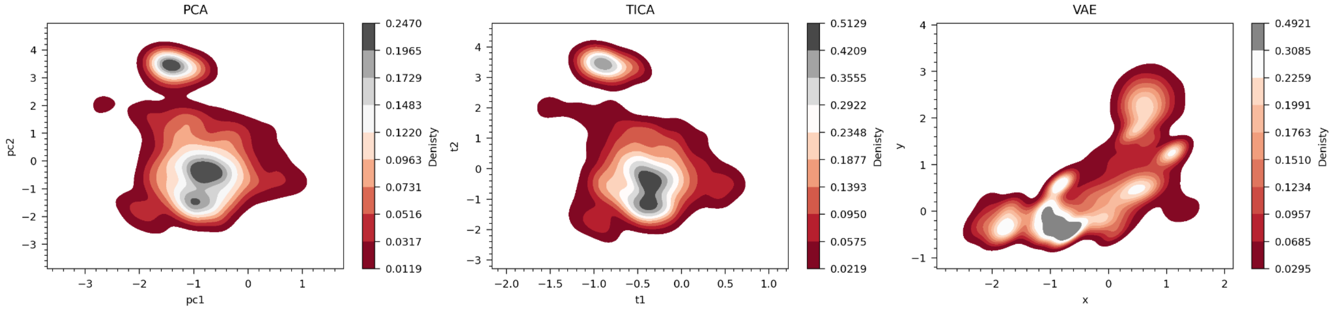


**Figure S6:** The projection of reduced representation of dihedral angles of Poly(U) is presented as (a) PCA, (b) TICA, and (c) VAE.

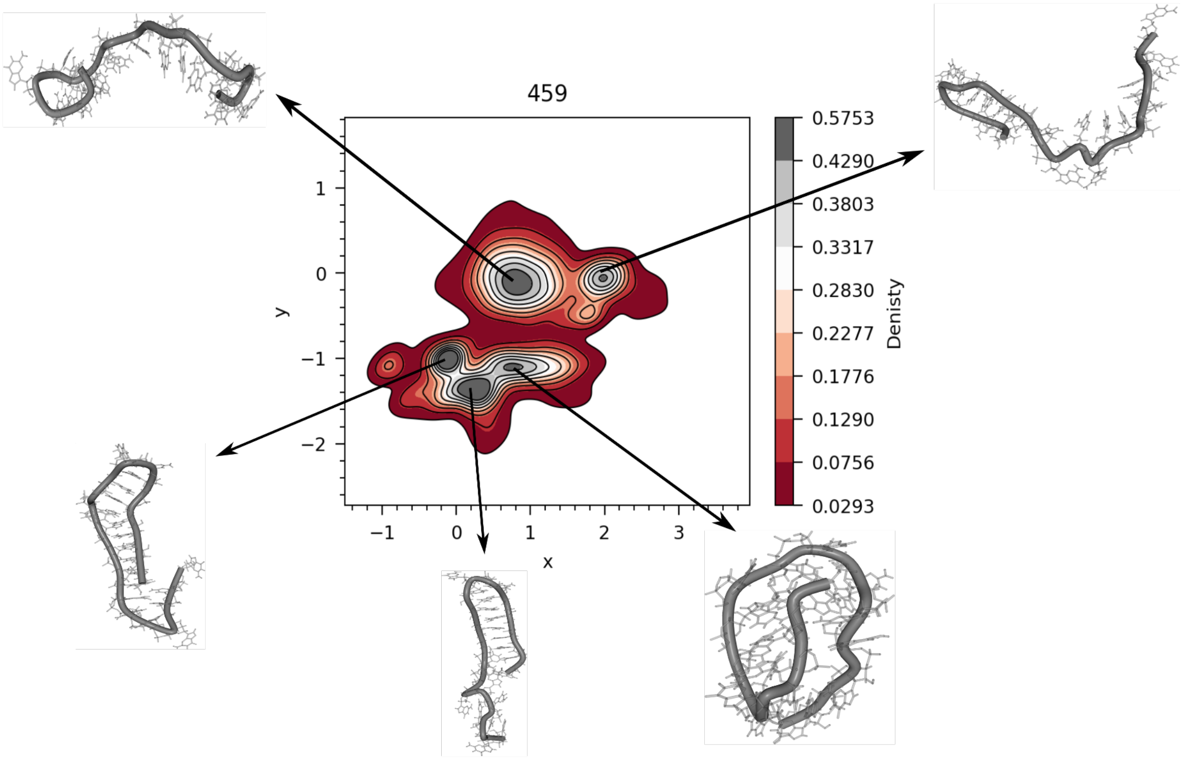


**Figure S7:** Structurally characterized energy landscape obtained by projection of reduced components of dihedral angle from VAE of  Poly(G) with four Major Structures at T ~ 459K.

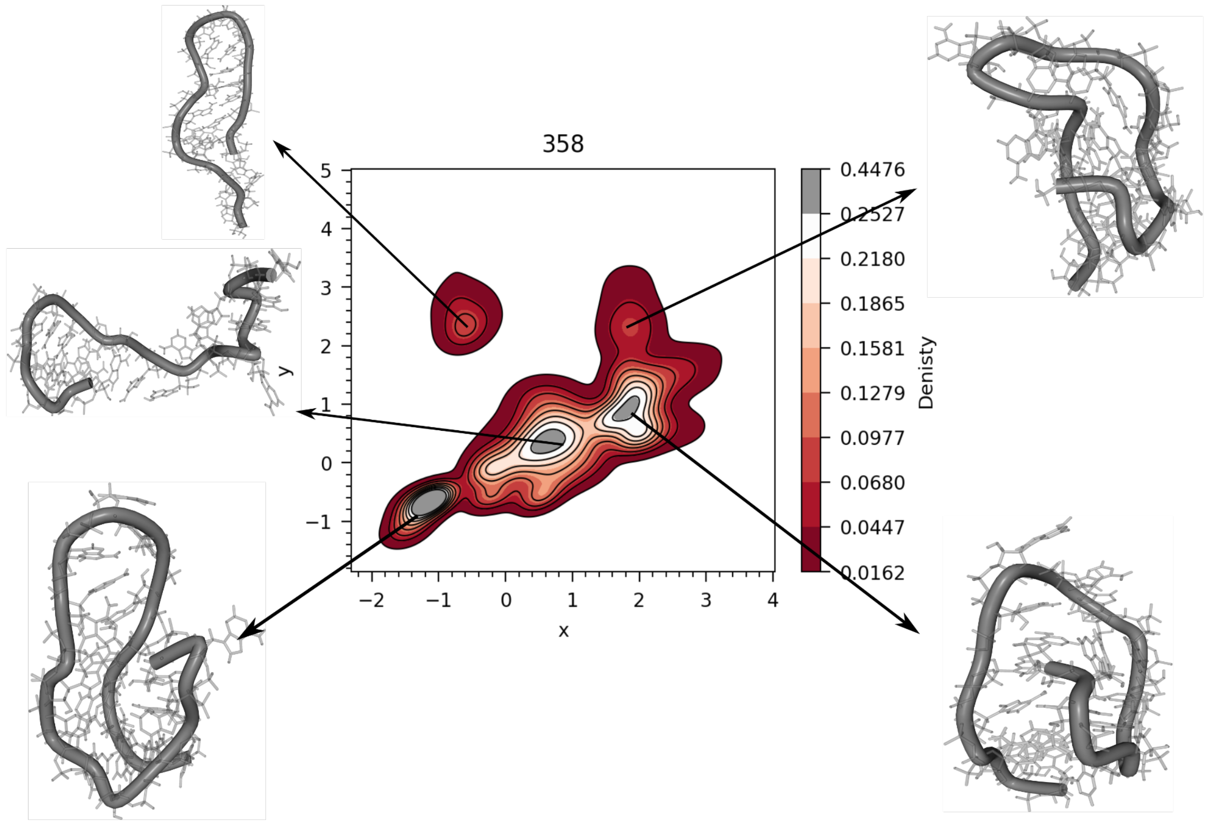


**Figure S8:** Structurally characterized energy landscape obtained by projection of reduced components of dihedral angle from VAE of Poly(A) with four Major Structures at T=358K.

### Poly(G)

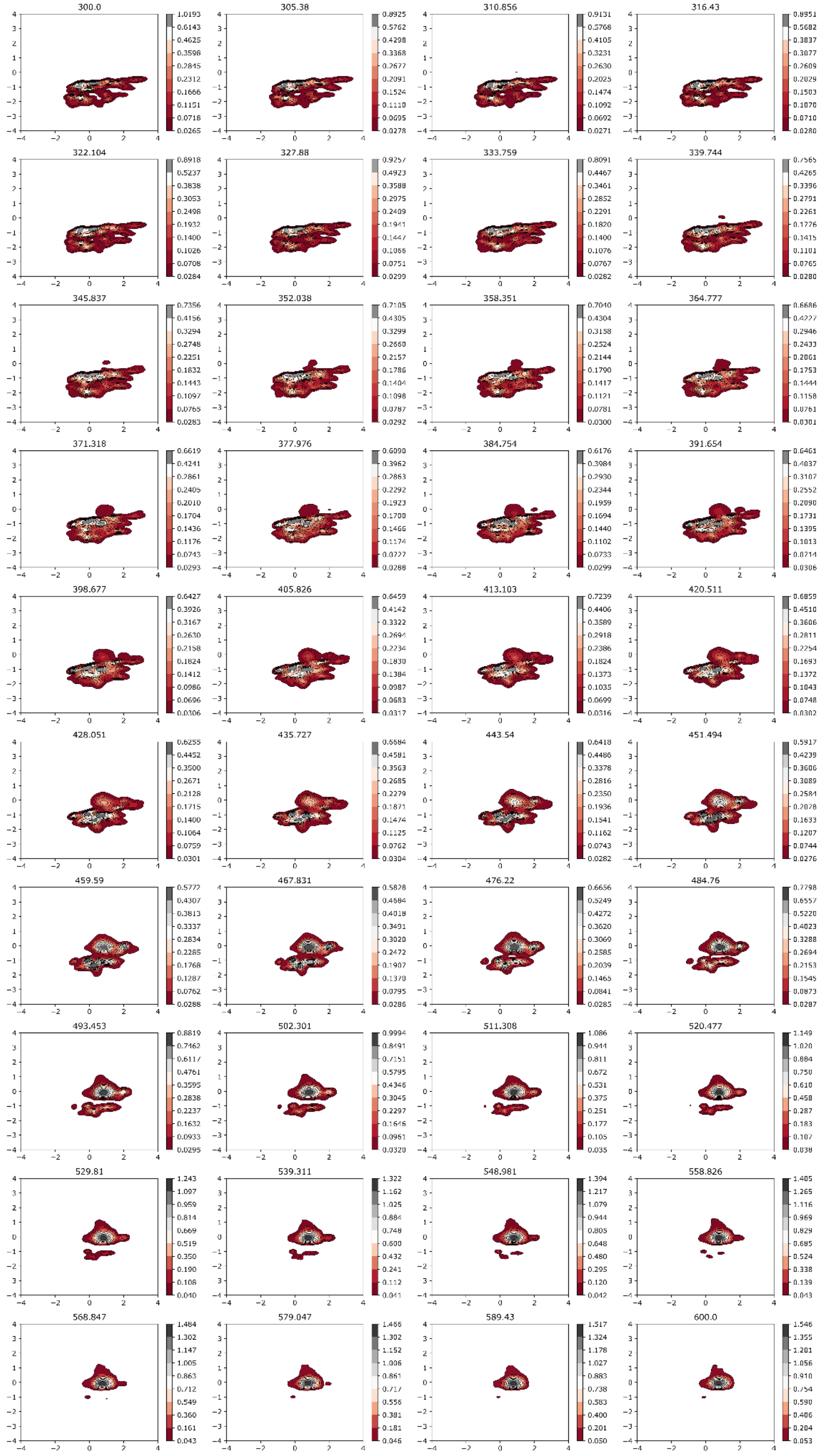


**Figure S9:** Temperature-Dependent Free Energy Landscapes Generated by the Latent Space of VAE for Guanine. The temperature of the landscape is written above the diagram.

### Poly(A)

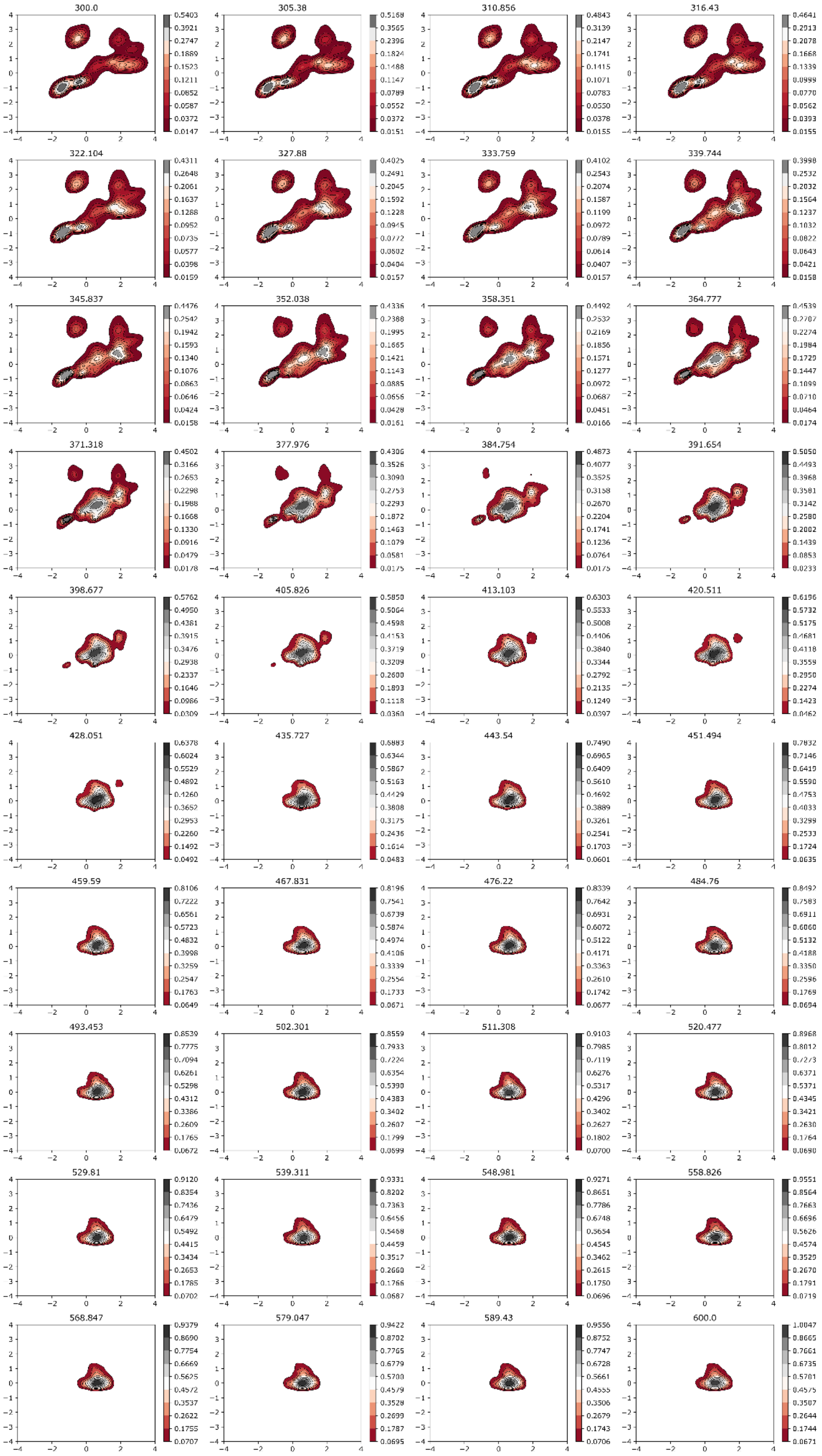

**Figure S10:**Temperature-Dependent Free Energy Landscapes Generated by the Latent Space of VAE for Adenine. The temperature of the landscape is written above the diagram.

#### Poly(C)

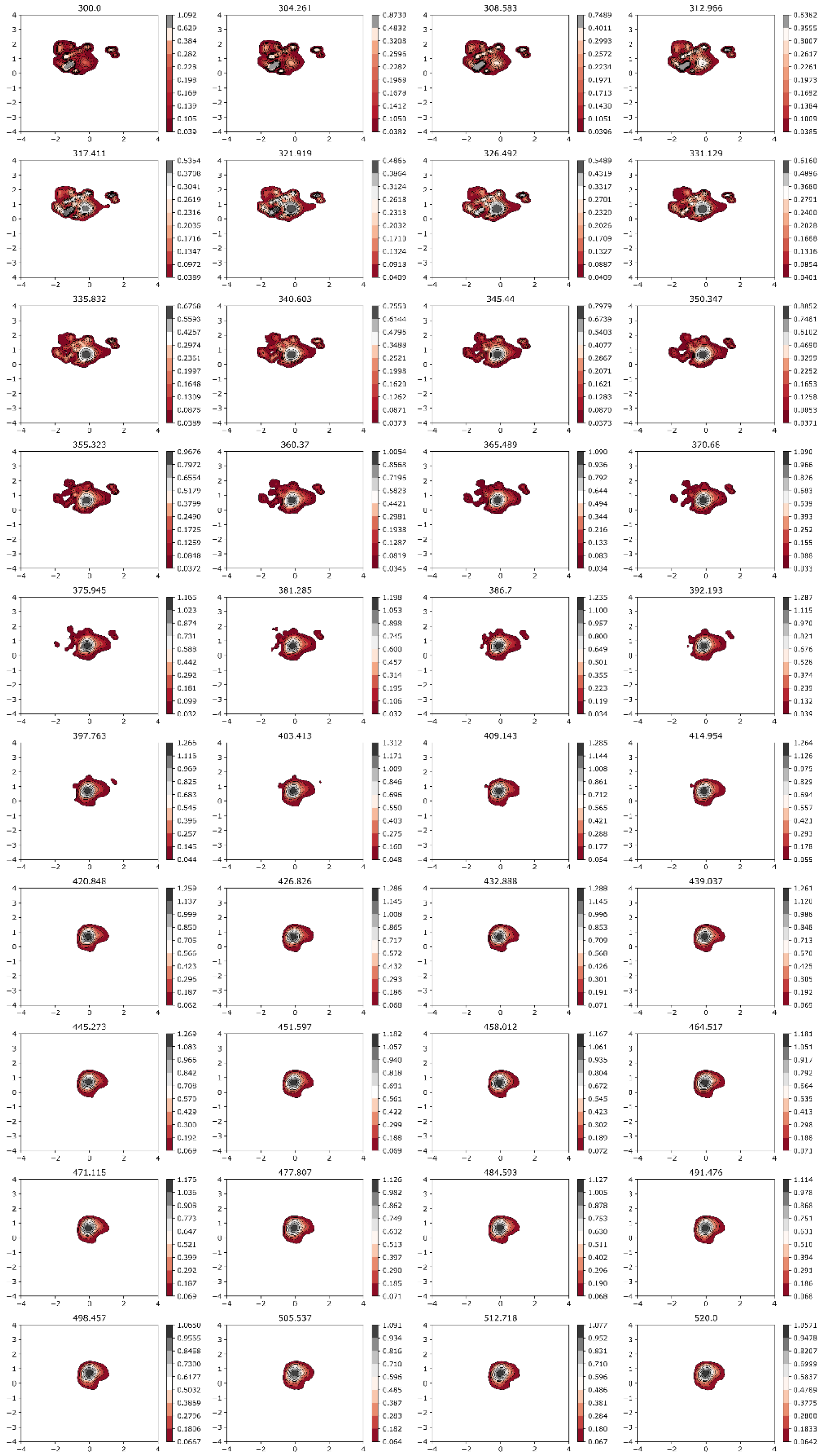

**Figure S11:** Temperature-Dependent Free Energy Landscapes Generated by the Latent Space of VAE for Cytosine. The temperature of the landscape is written above the diagram.

#### Poly(U)

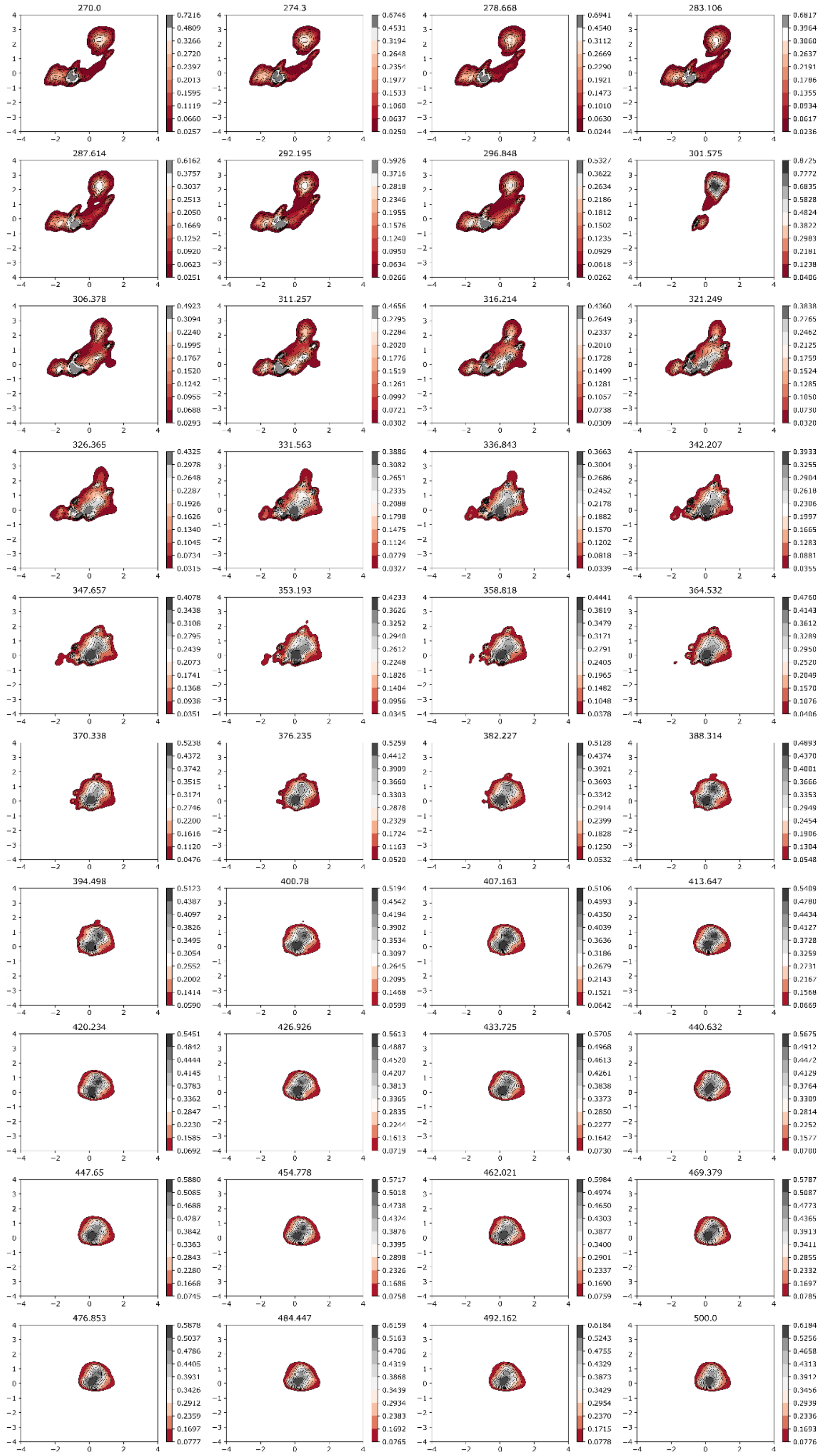

**Figure S12:** Temperature-Dependent Free Energy Landscapes Generated by the Latent Space of VAE for Cytosine. The temperature of the landscape is written above the diagram.

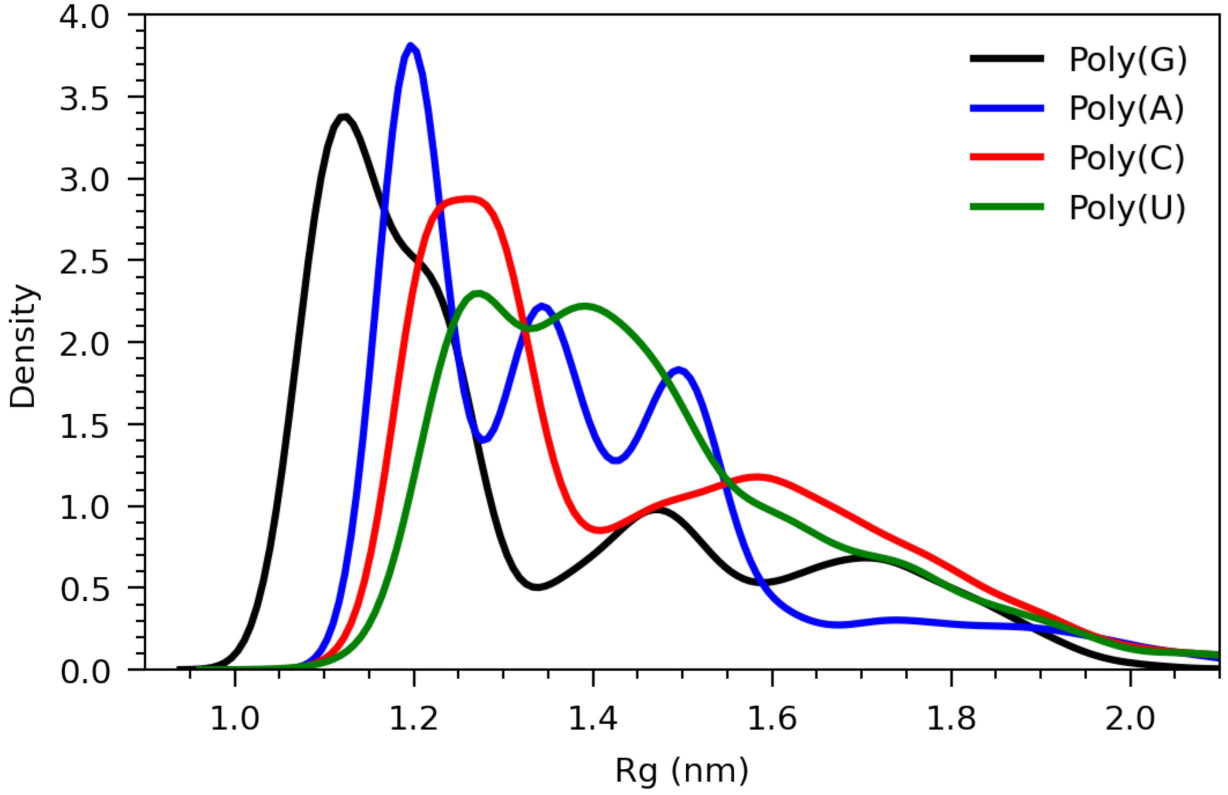


**Figure S13:** Distribution of Rg of Poly G at T=300K in black, Poly A at T = 300K in blue, Poly C at T=300K in red, and Poly U at T=306K in green.

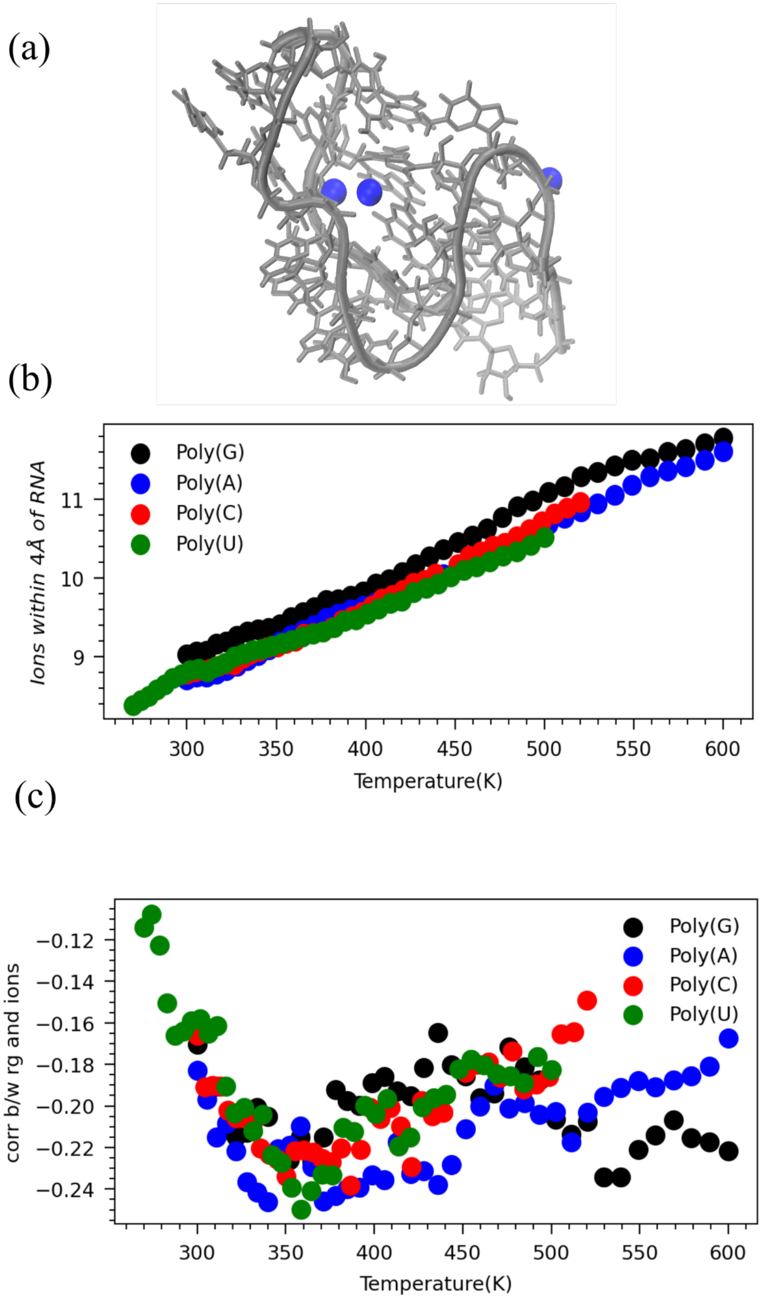


**Figure S14:** Dynamics of ions around RNA. (a) Illustration of $Na^+$ ions around Poly G structure shows ions help RNA to compact structure by overcoming coulombic repulsions. (b) The average number of ions within 4 Angstrom of RNA across temperatures. (c) correlation between Rg of RNA and number of $Na^+$ within 4 angstroms of RNA across temperature.

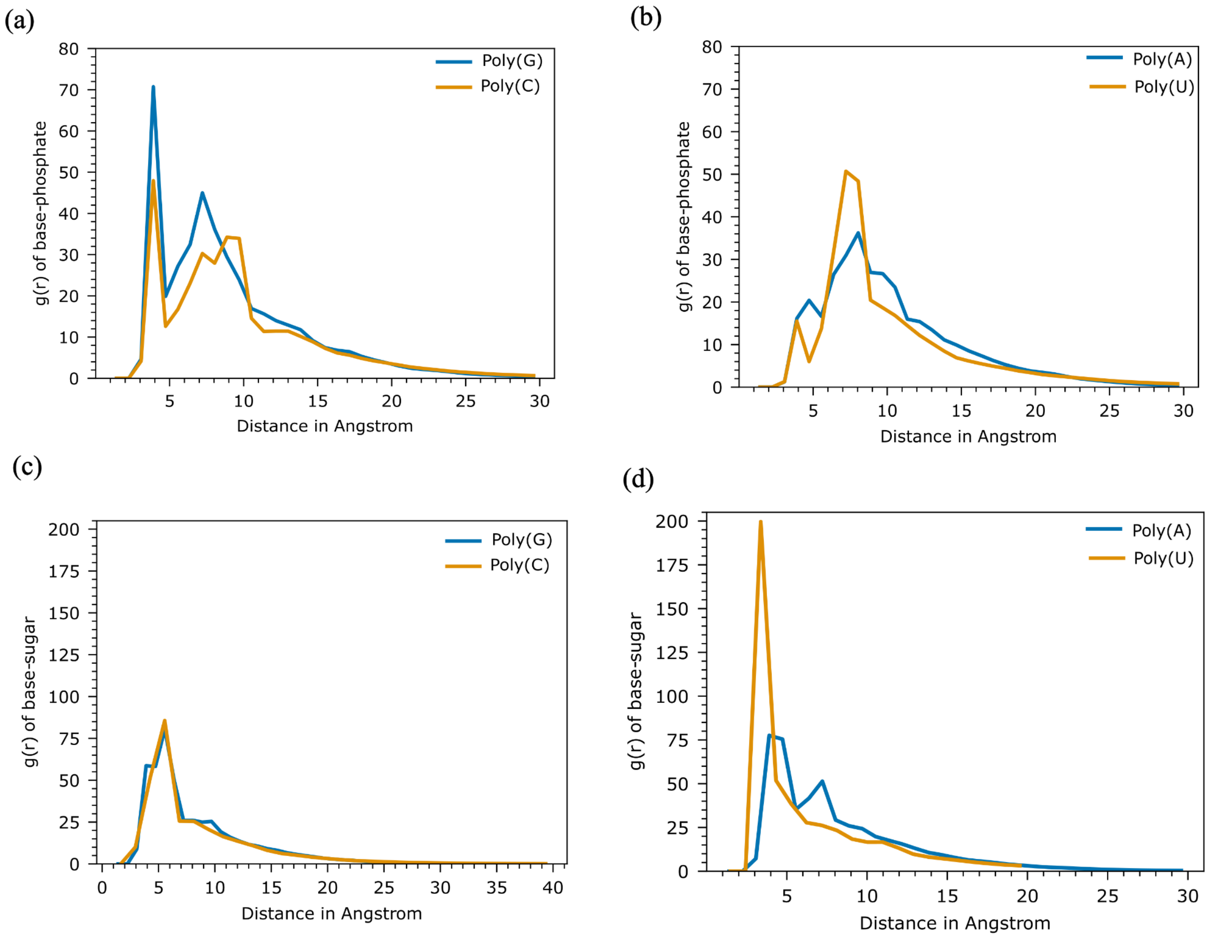


**Figure S15:** Studying the interaction through radial distribution function between (a) base-phosphate in poly G and Poly C, (b) base-phosphate in Poly A and Poly U, (c) base-sugar in poly G and poly C, and (d) base-sugar in Poly A and Poly U.

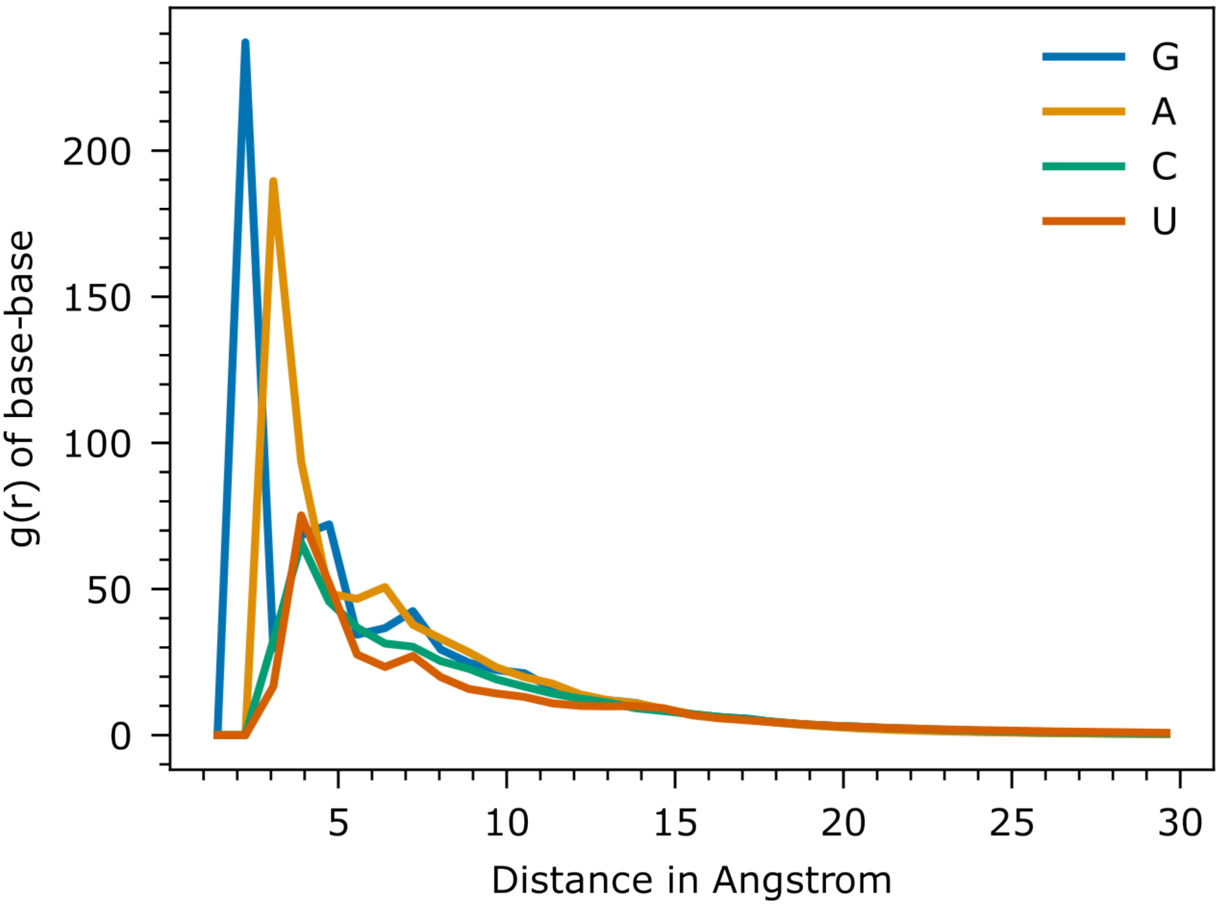


**Figure S16:** Radial distribution function between bases of Poly G, Poly A, Poly C, and Poly U.

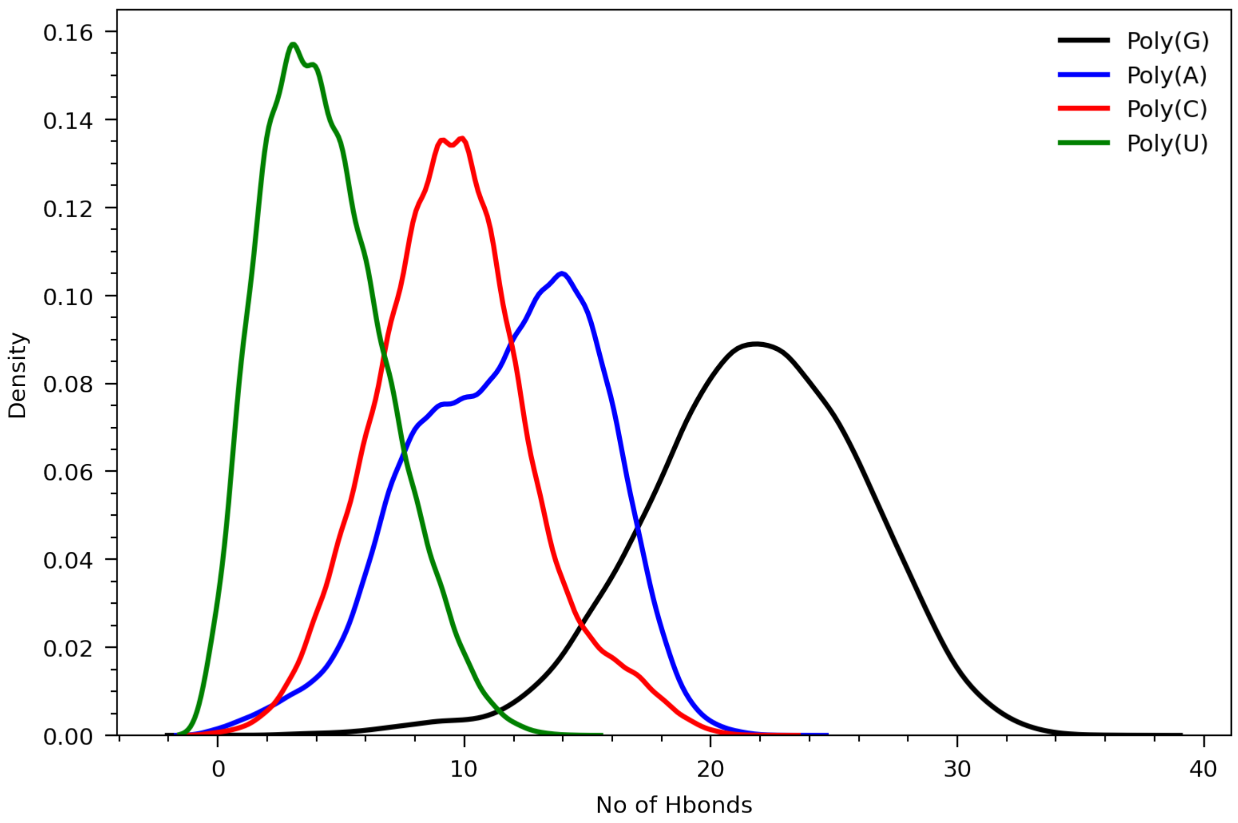


**Figure S17:** Distribution of hydrogen bond counts of Poly G at T=300K in black, Poly A at T = 300K in blue, Poly C at T=300K in red, and Poly U at T=306K in green.
